# The Effect of Transcranial Direct Current and Magnetic Stimulation on Fear Extinction and Return of Fear: A meta-analysis and Systematic Review

**DOI:** 10.1101/2023.09.11.557284

**Authors:** Grace L.T. Lei, Cora S.W. Lai, Tatia M.C. Lee, Charlene L.M. Lam

**Affiliations:** The State Key Laboratory of Brain and Cognitive Sciences, The University of Hong Kong, Hong Kong, China; School of Biomedical Sciences, LKS Faculty of Medicine, The University of Hong Kong, Pokfulam, Hong Kong; Laboratory of Neuropsychology and Human Neuroscience, The University of Hong Kong, Hong Kong, China; Laboratory of Clinical Psychology and Affective Neuroscience, The University of Hong Kong, Hong Kong, China; Guangdong-Hong Kong-Macao Greater Bay Area Center for Brain Science and Brain-Inspired Intelligence, Guangzhou, China

**Keywords:** transcranial direct current stimulation, tDCS, transcranial magnetic stimulation, TMS, fear conditioning, fear extinction, return of fear, fear retention

## Abstract

Anxiety and fear-related disorders are among the most prevalent mental illnesses. Non-invasive brain stimulation methods such as transcranial direct current stimulation (tDCS) and transcranial magnetic stimulation (TMS) have been employed to modulate anxiety and fear-related symptoms, but their therapeutic effects remain inconclusive. Pavlovian conditioning and extinction are experimental analogues of exposure therapy that investigate the neural mechanisms of fear extinction and return of fear. We conducted a meta-analysis and qualitative review on the effects of tDCS and TMS on fear extinction and return of fear in non-primate animals and humans. Results show that both anodal and cathodal tDCS over the prefrontal cortex inhibit short-term contextual and cued fear retrieval in animal models. In human studies, anodal tDCS over the medial/ventromedial prefrontal cortex enhances fear extinction, whereas TMS over the dorsolateral/ventromedial prefrontal cortex inhibits return of fear. Our findings suggest the optimal non-invasive brain stimulation protocols for threat extinction in humans.

## Introduction

Anxiety and fear-related disorders are among the most prevalent mental disorders (Bandelow & Michaelis, 2015). With 3.4% (264 million) of the global population a=:Jected (WHO, 2017) and a lifetime prevalence of 33.7% (Bandelow & Michaelis, 2015), anxiety and fear-related disorders have a significant impact on both individuals and society. More specifically, these disorders account for 24.6 million disability-adjusted life-years (WHO, 2017), lost productivity at work, reduced quality of life (Martino et al., 2019), a higher risk of mortality (van Hout et al., 2004), and a significant financial burden (Wittchen et al., 2011) for the affected individuals and society as a whole.

Fear conditioning paradigms are commonly used to study threat acquisition and its extinction in non-primate animals and humans. During the acquisition phase, an initially neutral stimulus (NS) is paired with an aversive unconditioned stimulus (US). After repeated pairings, the NS becomes a conditioned stimulus (CS), eliciting a conditioned response (CR) on its own. Cued fear conditioning and contextual fear conditioning are the two specific models used to measure threat learning and extinction under special circumstances. During cued fear conditioning, an NS (e.g., a tone) is paired with a US (e.g., an electrical foot shock). The NS then becomes a CS through repeated pairings, which triggers a CR (e.g., freezing) (VanElzakker et al., 2014). During contextual fear conditioning, a US is presented in a neutral context (e.g., a chamber), with subsequent re-exposure to that context triggering the CR on its own (Maren et al., 2013).

Extinction refers to a decrease in the conditioned fear response to a CS. It occurs when the CS-US association is gradually extinguished by repeatedly presenting the CS in the absence of the aversive US. During extinction learning, a newly formed association (CS-no US) is thought to compete with and inhibit the original fear association (CS-US), resulting in a reduction of the CR (Bouton, 2004). A maladaptive extinction learning process may contribute to the development and maintenance of anxiety and fear-related disorders (Tortella-Feliu et al., 2019).

The neural circuits of fear extinction and retention in non-primate animals involve the amygdala, hippocampus, and prefrontal cortex (PFC) (Herry et al., 2010; Moustafa et al., 2013; Quirk & Mueller, 2008). The sensory information of the CS and US is projected to the basolateral complex (BLA) of the amygdala and then to the central nucleus (CEA) (VanElzakker et al., 2014). Distinct neurons in the BLA respond differentially to fear conditioning and extinction processes (Herry et al., 2008). The firing rates of certain BLA neurons increase when rodents are exposed to a CS in the extinction context after extinction learning (Hobin et al., 2003). Moreover, local inhibition in BLA increases during expression of fear extinction (Ehrlich et al., 2009). In addition to the BLA, the CEA also receives projections from intercalated (ITC) cells. ITC neurons are crucial in fear extinction (Mańko et al., 2011), with findings revealing ITC cell lesions lead to deficits in fear extinction (Likhtik et al., 2008). Additionally, stimulus- and context-specific changes in neuronal responses to conditioned stimuli are observed in the lateral (CEl) and medial (CEm) CEA during fear extinction (Whittle et al., 2021). Whilst the amygdala is essential in cued fear extinction, the hippocampus plays a vital role in the extinction and retention of contextual fear (Moustafa et al., 2013). The hippocampus is crucial for representing and remembering context (Holland & Bouton, 1999), and hippocampal lesions are found to impair contextual fear extinction (Ji & Maren, 2007). In addition, pharmacological inactivation of the hippocampus produces deficits in the retrieval of contextual fear (Zelikowsky et al., 2012).

The infralimbic cortex (IL) projects to the ITC of the amygdala, which inhibits the outputs of the BLA and CEA, thereby suppressing fear expression (Berretta et al., 2005; Ehrlich et al., 2009). Basal amygdala neurons targeting the infralimbic (IL) subdivision exhibit cell-type-specific plasticity during fear extinction (Senn et al., 2014). During extinction learning, IL cells are activated in response to the presentation of a conditioned context (Thompson et al., 2010). Moreover, IL cells have been shown to respond to CS presentation during a retention test (Milad & Quirk, 2002). Studies have also revealed a negative correlation between the activities of IL neurons and the retention of fear memory (Burgos-Robles et al., 2007; Herry & Garcia, 2002). Infralimbic lesions do not interfere with fear conditioning or extinction, but enhance the return of fear 24 hours later (Lebrón et al., 2004; Quirk et al., 2000).

Studies examining the neural circuits of fear conditioning in humans have reported similar findings to studies using animal models, with the amygdala, hippocampus, ventromedial PFC (vmPFC), and dorsolateral PFC (dlPFC) (Etkin et al., 2011; Quirk & Mueller, 2008) implicated in both human and animal models. Several parts of the amygdala are thought to be crucial for the extinction of fear. Specifically, signals of the US and CS converge and are processed in the BLA, which then sends its output to the CEA (Barad et al., 2006). Neurons in the CEA initiate the physiological and behavioural response to threat (Sah & Westbrook, 2008). ITC amygdala neurons receive signals of the CS from the BLA and send inhibitory projections to the CEA to inhibit the expression of fear (Likhtik et al., 2008).

As well as the amygdala, the dlPFC, and vmPFC are also implicated in fear extinction (Phelps et al., 2004). The dlPFC is essential to the consolidation, retrieval, and reactivation of memory traces (Sandrini et al., 2013; Simons & Spiers, 2003), and is likely to be involved in the reduction of the threat response and modulation of threat memory (Mungee et al., 2014). Cooperating with the amygdala, the vmPFC integrates information from multiple inputs to exert top-down control on the outputs of fear conditioning (Milad et al., 2007). Lesions of the vmPFC have been found to increase the right amygdala’s reactivity to aversive stimuli (Motzkin et al., 2015). Furthermore, the vmPFC is a crucial target of the hippocampus, which activates or inhibits fear expression according to the context of acquisition (VanElzakker et al., 2014). The vmPFC-hippocampal network thus plays a vital role in context-dependent fear extinction recall (Kalisch et al., 2006). Prior studies on the neural circuits related to fear extinction shed lights on the possibility of enhancing extinction by stimulating the relevant brain regions, among which the PFC is a common targeting area.

Advances in neuroscience methods have contributed to the identification of the neural mechanisms of threat memory and the development of new treatment protocols. Non-invasive brain stimulation (NIBS), including transcranial direct current stimulation (tDCS) and transcranial magnetic stimulation (TMS), has demonstrated a promising therapeutic effect on both anxiety disorders (Vicario, Salehinejad, et al., 2020) and fear memory (Borgomaneri et al., 2021). tDCS alters brain functions with direct electrical currents that pass through the cerebral cortex (Nitsche & Paulus, 2000). Part of the current is absorbed by the skull, whereas other parts penetrate the scalp and modulate cortical excitability (Reed & Cohen Kadosh, 2018). Typically, electrical currents (1 - 2 mA) are delivered via two or more electrodes of opposite polarities (i.e., anode and cathode) placed on the scalp, which increases or decreases neuronal activity by modulating the membrane potential of the stimulated neurons (Nitsche & Paulus, 2000). Anodal stimulation depolarizes the neurons and increases cortical excitability, whereas cathodal stimulation hyperpolarizes neurons and inhibits the occurrence of action potentials (Nitsche et al., 2008).

Unlike tDCS, TMS employs magnetic fields to induce electrical discharges of target areas of the brain. Trains of magnetic pulses are delivered via a coil positioned on the scalp. When the magnetic pulses are repetitively applied, they can modulate cortical excitability. Repetitive TMS (rTMS) can modify brain function for minutes to hours (Huang et al., 2005; Iyer et al., 2003). In general, low-frequency stimulation (≤1 Hz) has inhibitory e=:Jects, whereas high-frequency stimulation (≥5 Hz) has excitatory e=:Jects (Klomjai et al., 2015).

Given the established knowledge of the neural circuits involved in fear extinction, NIBS has been employed to facilitate the extinction of threat responses in non-primate animals and human studies. Previous studies on the treatment effect of tDCS and TMS on anxiety and fear-related disorders have reported mixed findings. Some studies have suggested that tDCS and TMS reduced the symptoms of anxiety and fear-related disorders, whereas others have observed no significant improvement (see Cirillo et al., 2019 and Stein et al., 2020 for a review). The fear extinction model is commonly used to study the psychopathology of anxiety disorders and their neural mechanisms (VanElzakker et al., 2014). Exploring the effects of tDCS and TMS on fear extinction and the return of fear can contribute to the development of interventions combining NIBS and exposure therapy. Here, we reviewed studies exploring the effects of tDCS and TMS on threat extinction in non-primate animals and humans with the goals of synthesising the current state of knowledge and shedding light on the development of effective NIBS treatment protocols. Most of the relevant studies employed the frontal cortex or PFC as the targeting brain region.

## Transparency and Openness

Data and codes of this study are available online (https://osf.io/hd7gv/?view_only=5e9379ef6b274841ac7242e278104e3a).

## Method

### Literature searching and selection

Following the guidelines of Preferred Reporting Items for Systematic Reviews and Meta-Analyses (Page et al., 2021), we searched PubMed, Web of Science, PsycINFO, and Cochrane Library using the following search strategies: 1) (“transcranial direct current stimulation” OR “tDCS”) [Title/Abstract] AND (“fear conditioning” OR “fear extinction” OR “fear memory” OR “return of fear” OR “fear retention” OR “fear recall” OR “threat” OR “threat memory” OR “Pavlovian conditioning”) [Title/Abstract]; 2) (“transcranial electrical stimulation” OR “tES”) [Title/Abstract] AND (“fear conditioning” OR “fear extinction” OR “fear memory” OR “return of fear” OR “fear retention” OR “fear recall” OR “threat” OR “threat memory” OR “Pavlovian conditioning”) [Title/Abstract]; 3) (“transcranial magnetic stimulation” OR “TMS”) [Title/Abstract] AND (“fear conditioning” OR “fear extinction” OR “fear memory” OR “return of fear” OR “fear retention” OR “fear recall” OR “threat” OR “threat memory” OR “Pavlovian conditioning”) [Title/Abstract]. Additionally, we searched Google Scholar with the following keywords: “tDCS and fear conditioning”, “tES and fear conditioning”, as well as “TMS and fear conditioning”. The date range is extended up to 30 Sept 2022. We also screened the reference lists of published systematic reviews of the effect of tDCS and TMS on fear conditioning (Borgomaneri et al., 2021; Herrmann et al., 2019; Marković et al., 2021; Yosephi et al., 2019).

The detailed literature selection procedures are illustrated in Figure 1. Inclusion criteria were as follows: randomized and sham-controlled trials published in English, animal or human subjects, trials concerning fear conditioning, and trials employed tDCS or TMS methods. The exclusion criteria were as follows: non-controlled or non-randomized trials, case reports or case series, reviews or commentaries, trials of paradigm other than the fear conditioning model, trials of interventions other than tDCS or TMS, trials without a control sham group, and duplicated data sets. Finally, we identified eight articles on the tDCS effect and three articles on the TMS effect on fear conditioning among animals. For human studies, we included eight articles on the tDCS effect and three articles on the TMS effect. Altogether, there were 246 animal subjects in the active tDCS group and 137 subjects in the sham group, whereas 45 subjects in the active TMS group and 43 subjects in the sham group. In human tDCS studies, the active group consisted of 236 participants and the sham group comprised 156 participants. In human TMS studies, 102 participants were in the active group and 95 participants in the sham group. Detailed information on the included studies is listed in Tables 1 and 2.

**Figure 1.**
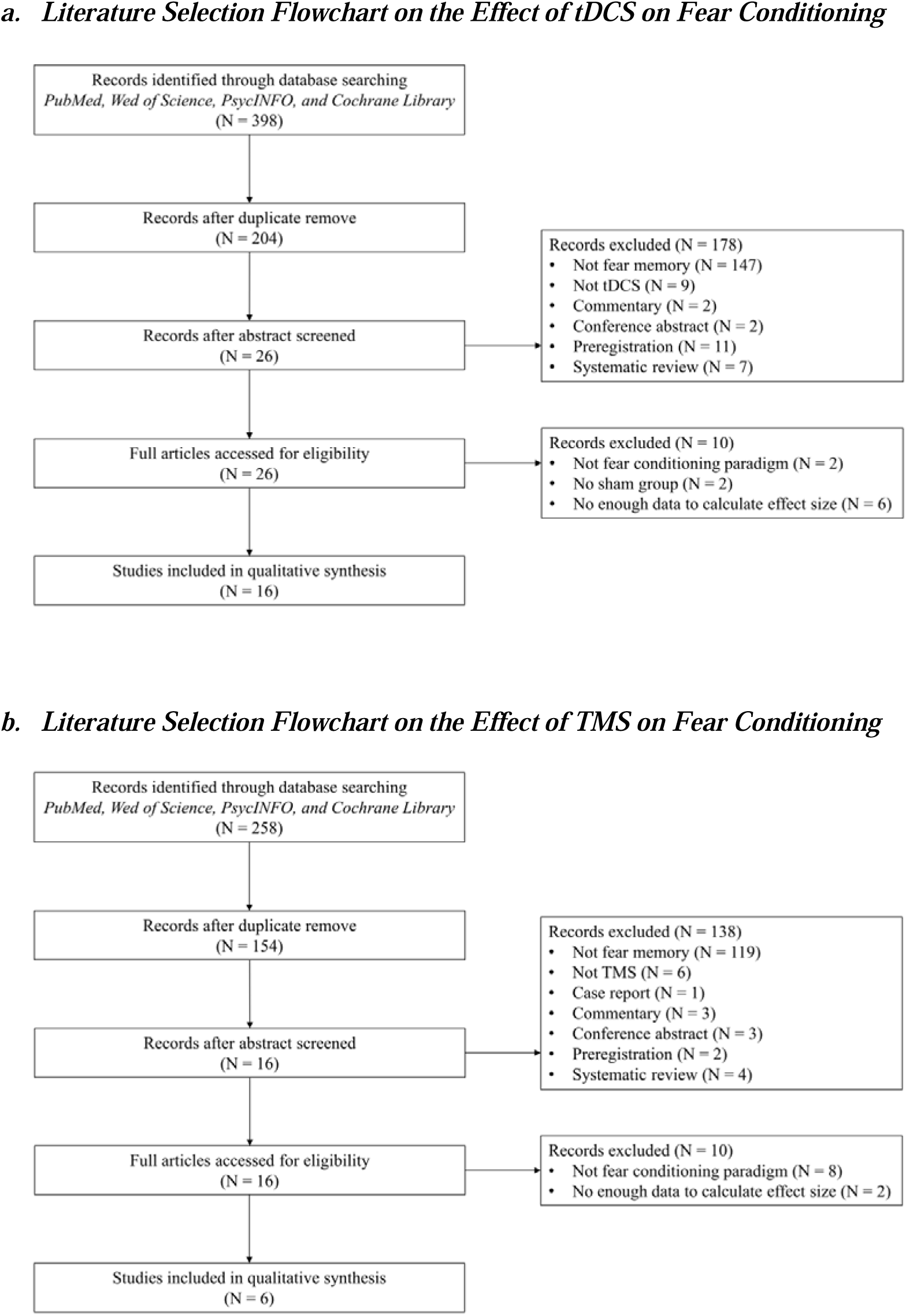

**Table 1a.**
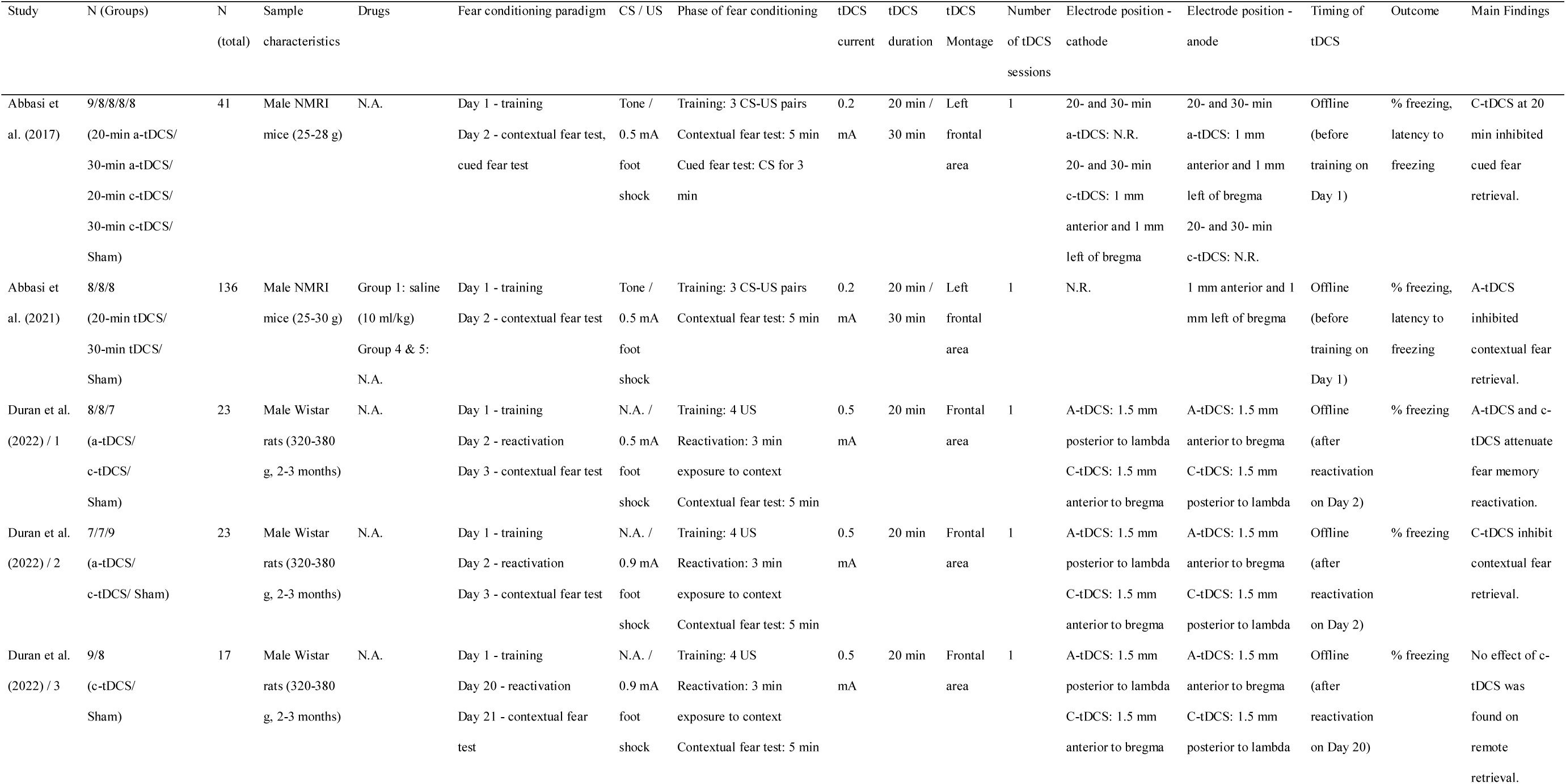
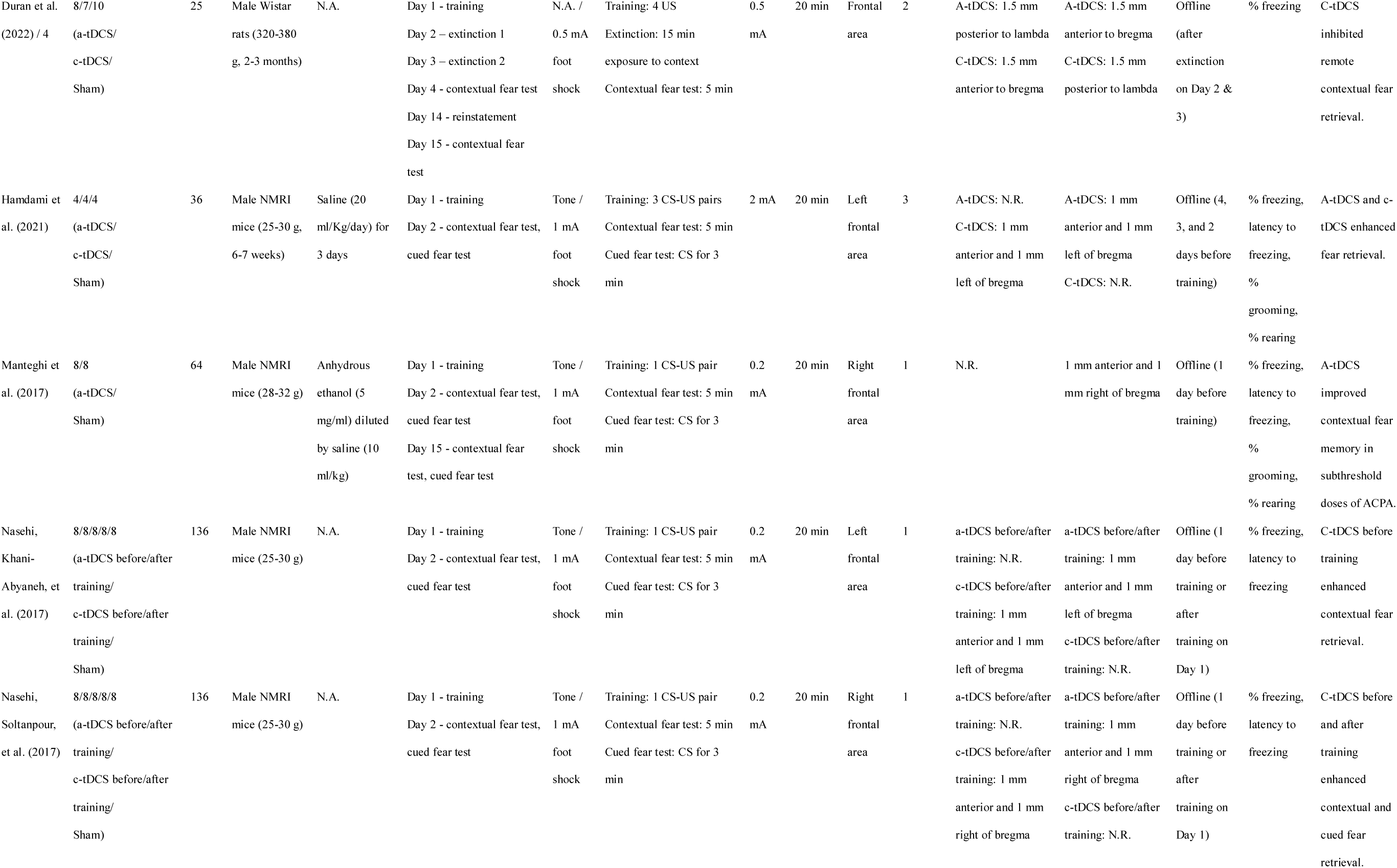
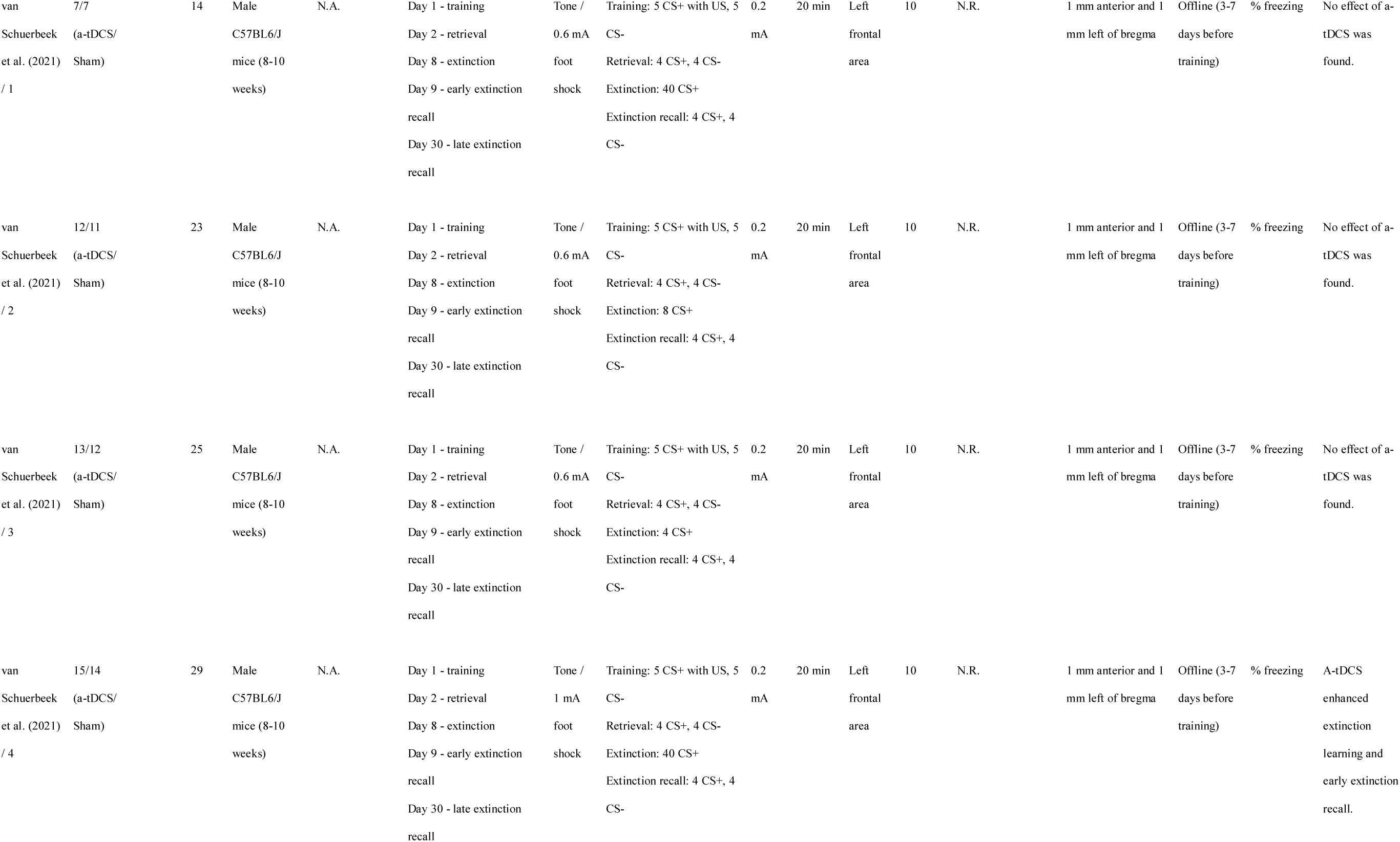
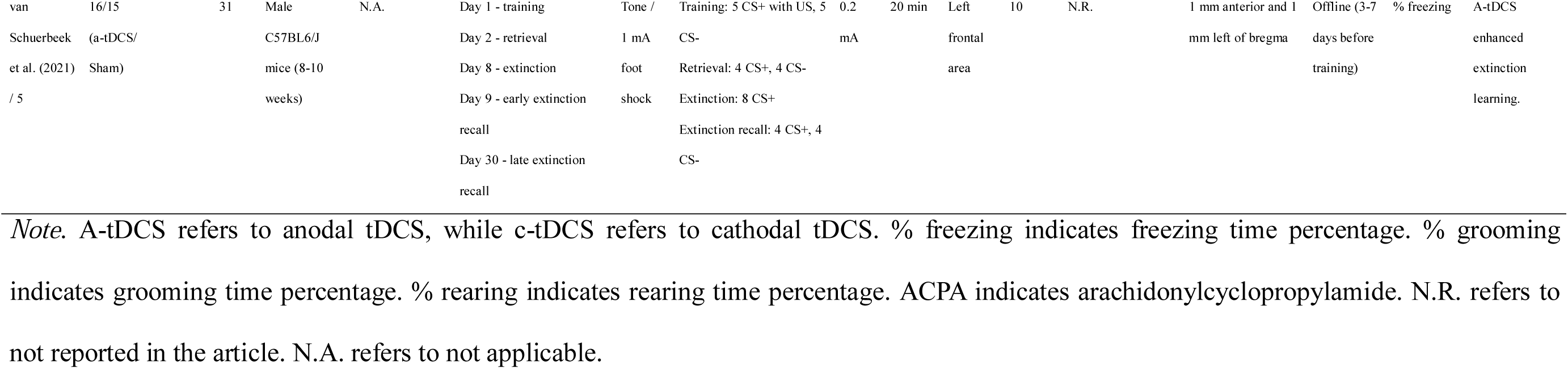
Summary of the Included Animal Studies on the effect of tDCS on Fear Conditioning.

**Table 1b.**
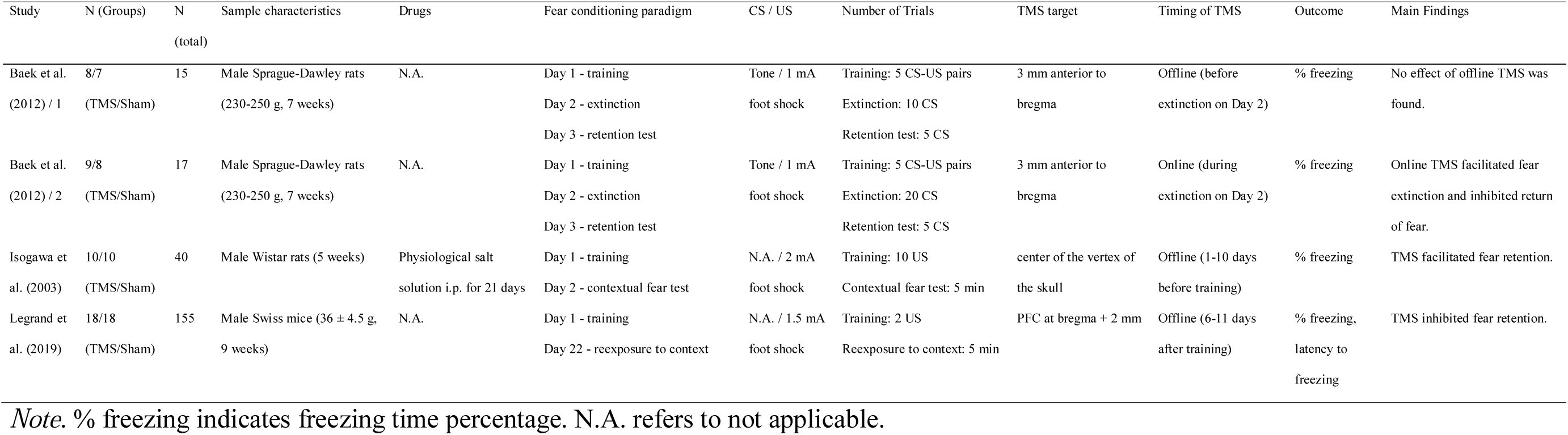
Summary of the Included Animal Studies on TMS and Fear Conditioning.

| Study | N (Groups) | N<br>(total) | Sample characteristics | Drugs | Fear conditioning paradigm | CS / US | Number of Trials | TMS target | Timing of TMS | Outcome | Main Findings |
| --- | --- | --- | --- | --- | --- | --- | --- | --- | --- | --- | --- |
| Baek et al.<br>(2012) / 1 | 8/7<br>(TMS/Sham) | 15 | Male Sprague-Dawley rats<br>(230-250 g, 7 weeks) | N.A. | Day 1 - training<br>Day 2 - extinction<br>Day 3 - retention test | Tone / 1 mA<br>foot shock | Training: 5 CS-US pairs<br>Extinction: 10 CS<br>Retention test: 5 CS | 3 mm anterior to<br>bregma | Offline (before<br>extinction on Day 2) | % freezing | No effect of offline TMS was<br>found. |
| Baek et al.<br>(2012) / 2 | 9/8<br>(TMS/Sham) | 17 | Male Sprague-Dawley rats<br>(230-250 g, 7 weeks) | N.A. | Day 1 - training<br>Day 2 - extinction<br>Day 3 - retention test | Tone / 1 mA<br>foot shock | Training: 5 CS-US pairs<br>Extinction: 20 CS<br>Retention test: 5 CS | 3 mm anterior to<br>bregma | Online (during<br>extinction on Day 2) | % freezing | Online TMS facilitated fear<br>extinction and inhibited return<br>of fear. |
| Isogawa et<br>al. (2003) | 10/10<br>(TMS/Sham) | 40 | Male Wistar rats (5 weeks) | Physiological salt<br>solution i.p. for 21 days | Day 1 - training<br>Day 2 - contextual fear test | N.A. / 2 mA<br>foot shock | Training: 10 US<br>Contextual fear test: 5 min | center of the vertex of<br>the skull | Offline (1-10 days<br>before training) | % freezing | TMS facilitated fear retention. |
| Legrand et<br>al. (2019) | 18/18<br>(TMS/Sham) | 155 | Male Swiss mice (36 ± 4.5 g,<br>9 weeks) | N.A. | Day 1 - training<br>Day 22 - reexposure to context | N.A. / 1.5 mA<br>foot shock | Training: 2 US<br>Reexposure to context: 5 min | PFC at bregma + 2 mm | Offline (6-11 days<br>after training) | % freezing,<br>latency to<br>freezing | TMS inhibited fear retention. |

*Note.* % freezing indicates freezing time percentage. N.A. refers to not applicable.

**Table 2a.**
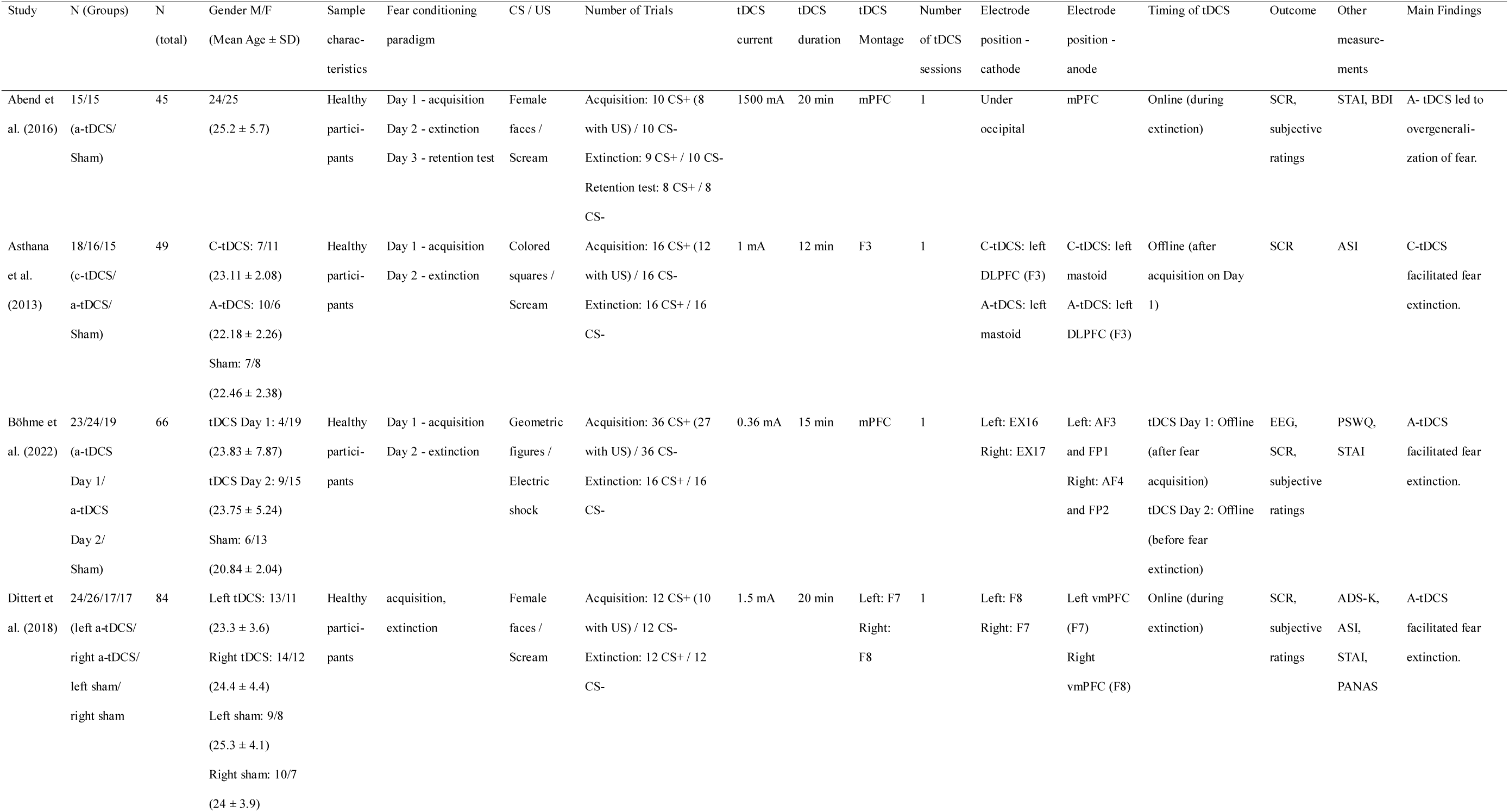
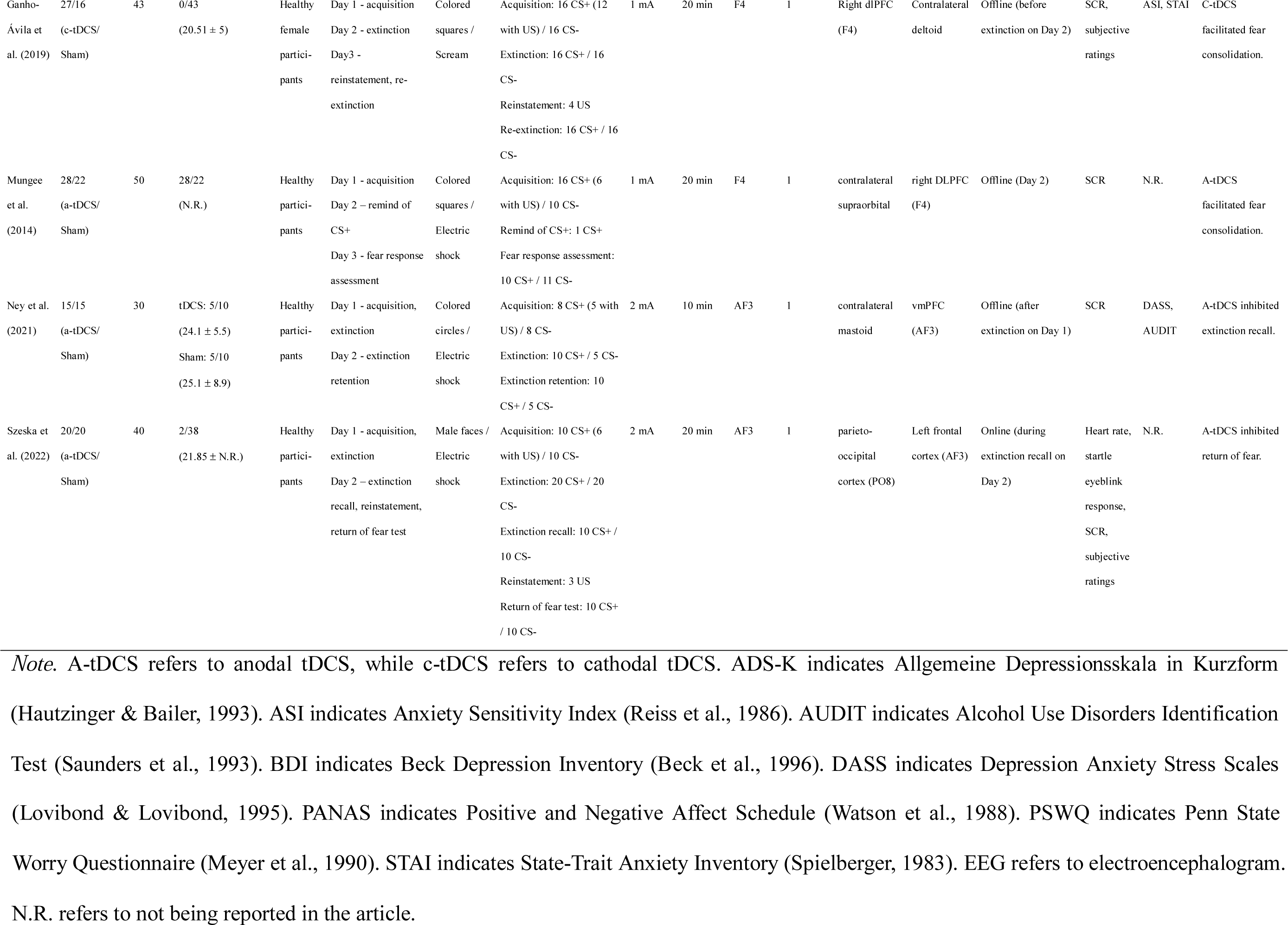
Summary of the Included Human Studies on the effect of tDCS on Fear Conditioning.

**Table 2b.** Summary of the Included Human Studies on TMS and Fear Conditioning.

| Study | N (Groups) | N<br>(total) | Gender M/F<br>(Mean Age ± SD) | Sample<br>characteristics | Fear conditioning paradigm | CS / US | Number of Trials | TMS target | Timing of TMS | Outcome | Other<br>measurements | Main Findings |
| --- | --- | --- | --- | --- | --- | --- | --- | --- | --- | --- | --- | --- |
| Borgomaneri<br>et al. (2020) | 14/14/14<br>(left TMS/<br>right TMS/<br>Sham) | 84 | Left TMS: 6/8<br>(23.9 ± 2.3)<br>Right TMS: 5/9<br>(23.1 ± 2.6)<br>Sham: 6/8<br>(23.2 ± 1.8) | Healthy<br>participants | Day 1 - acquisition<br>Day 2 - reactivation<br>Day 3 - recall, extinction,<br>reinstatement test | Visual scenes /<br>Electric shock | Acquisition: 16 CS+ (10 with US) / 16 CS-<br>Reactivation: 2 CS+<br>Recall: 4 CS+ / 4 CS-<br>Extinction: 12 CS+ / 12 CS-<br>Reinstatement: 3 US / 4 CS+ / 4 CS- | Left: left dlPFC<br>Right: right<br>dlPFC | Offline (after<br>reactivation on Day<br>2) | SCR, subjective<br>ratings | STAI, HADS | TMS on left and right<br>dlPFC inhibited return of<br>fear. |
| Deng et al.<br>(2021) / 1 | 16/19<br>(TMS/Sham) | 35 | TMS: 12/4<br>(23.31 ± 1.21)<br>Sham: 12/7<br>(23.05 ± 1.04) | Healthy<br>participants | Day 1 - acquisition<br>Day 2 - extinction<br>Day 3 - spontaneous recovery<br>test, reinstatement test | Colored squares /<br>Electric shock | Acquisition: 24 CS+ (12 with US) / 12 CS-<br>Extinction: 15 CS+ / 15 CS-<br>Spontaneous recovery: 15 CS+ / 15 CS-<br>Reinstatement: 3 US / 15 CS+ / 15 CS- | Left dlPFC (F3) | Offline (before<br>extinction on Day 2) | SCR | SDS, SAS,<br>MoCA, digit<br>span test | TMS on left dlPFC<br>inhibited return of fear. |
| Deng et al.<br>(2021) / 2 | 18/17<br>(TMS/Sham) | 35 | TMS: 9/9<br>(22 ± 0.52)<br>Sham: 6/11<br>(21.59 ± 0.59) | Healthy<br>participants | Day 1 - acquisition<br>Day 2 - extinction<br>Day 3 - spontaneous recovery<br>test, reinstatement test | Colored squares /<br>Electric shock | Acquisition: 24 CS+ (12 with US) / 12 CS-<br>Extinction: 15 CS+ / 15 CS-<br>Spontaneous recovery: 15 CS+ / 15 CS-<br>Reinstatement: 3 US / 15 CS+ / 15 CS- | Left dlPFC (F3) | Offline (after<br>extinction on Day 2) | SCR | SDS, SAS,<br>MoCA, digit<br>span test | TMS on left dlPFC<br>inhibited return of fear. |
| Guhn et al.<br>(2014) | 40/45<br>(TMS/Sham) | 85 | TMS: 21/19<br>(23.9 ± 3)<br>Sham: 22/23<br>(24.6 ± 4.5) | Healthy<br>participants | Day 1 - acquisition, extinction<br>learning<br>Day 2 - extinction recall | Male faces /<br>Scream | Acquisition: 16 CS+ (8 with US) / 16 CS-<br>Extinction learning: 20 CS+ / 20 CS-<br>Extinction recall: 20 CS+ / 20 CS- | mPFC | Offline (between<br>acquisition and<br>extinction learning<br>on Day 1) | EMG, fNIRS,<br>SCR, subjective<br>ratings | STAI, PANAS | TMS on mPFC facilitated<br>extinction learning and<br>inhibited return of fear. |

*Note.* EMG refers to electromyography, while fNIRS refers to functional near-infrared spectroscopy. HADS indicates Hospital Anxiety and Depression Scale (Zigmond & Snaith, 1983). MoCA indicates Montreal Cognitive Assessment (Nasreddine et al., 2005). PANAS indicates Positive and Negative Affect Schedule (Watson et al., 1988). SAS indicates Self-Rating Anxiety Scale (Zung, 1971). SDS indicates Self-Rating Depression Scale (Zung, 1965). STAI indicates State-Trait Anxiety Inventory (Spielberger, 1983). N.R. refers to not being reported in the article.

Additionally, some articles failed to meet the inclusion criteria due to different outcome measurements and inadequate data for calculating effect sizes. However, these studies also investigate the effect of tDCS or TMS on fear conditioning and extinction. Main findings of these articles are listed in Table S1.

### Primary outcome and data extraction

Freezing time percentage and latency to freezing are the primary efficacy outcomes for animal studies. We extracted the freezing time percentage and latency to freezing of the active group and the sham group during the contextual fear and cued fear tests. For human studies, skin conductance response (SCR) is the main efficacy outcome. We extracted the average discriminative SCR (i.e., SCR to CS+ minus SCR to CS-) of the active group and the sham group during extinction and tests of return of fear (ROF), respectively.

The following information was also identified: sample size of active and sham groups, sample characteristics, drugs injected for animals, other measurements for human participants, fear conditioning paradigm, setting, and timing of tDCS/TMS.

### Meta-analysis

The effect of tDCS/TMS on fear conditioning was quantified by Hedges’ g, a measure of effect size recommended for meta-analysis (Lakens, 2013). Additionally, 95% confidential intervals (CIs) of Hedges’ g were calculated for each study. The effect of anodal and cathodal tDCS was analyzed separately because they have the opposite effect on brain activations. In addition, the opposite electrode was placed on a reference position instead of areas involved in the fear circuit (see Tables 1a and 2a for details), which prevented us from combining the two types of stimulation into one meta-analysis.

For animal studies, separate meta-analyses were conducted to examine the effects of anodal and cathodal tDCS on short-term and long-term retentions of contextual fear memory, the effects of anodal and cathodal tDCS on short-term and long-term retentions of cued fear memory. Effect sizes were calculated by two behavioural outcomes in animals by comparing: 1) the difference in freezing time percentage and 2) the latency to freezing during the contextual fear test or cued fear test between the active and sham groups. A positive g value on the freezing time percentage indicated more fear responses during the retention test in the active group than in the sham group. On the contrary, a positive g value on latency to freezing time percentage indicated a smaller fear response during the retention test in the active group than in the sham group.

For human studies, separate meta-analyses were conducted to examine the effects of anodal and cathodal tDCS on fear extinction, the effects of anodal and cathodal tDCS on ROF, as well as the effects of TMS on fear extinction and ROF, respectively. Effect sizes of tDCS/TMS on fear extinction were calculated by comparing the difference of discriminative SCR or discriminative subjective ratings during the extinction phase between the active group and the sham group. A positive g value indicated a larger differentiation between CS+ and CS-during fear extinction in the active group than in the sham group. Similarly, effect sizes of tDCS/TMS on ROF were calculated by comparing the difference of discriminative SCR or discriminative subjective ratings during ROF test phase between the active and the sham group. A positive g value indicated a larger differentiation between CS+ and CS-during the ROF test in the active group than in the sham group.

The effect sizes were further processed with the “metafor" package in R following the template developed by (Yeung & Feldman, 2021). A random effect model was applied to analyze the overall effect size. Such a model assumes that each study estimated different values from the population parameters (Tufanaru et al., 2015). The assumption of heterogeneity was assessed with Cochran’s Q test, which was calculated as the weighted sum of squared differences between the effects of individual studies and the pooled effect across studies. Moreover, the I^2^ index was estimated to quantify the proportion of variance in the observed effect attributable to sampling error, with values of 25 %, 50 %, and 75 % reflecting small, medium, and large degrees of heterogeneity, respectively (Higgins & Thompson, 2002).

### Moderation analysis

Apart from the meta-analysis, moderation analysis was also conducted to test if the effect of tDCS/TMS on fear extinction and return of fear was moderated by several factors. The amplitude, duration, and targeting areas of tDCS were employed as moderators on the effect of tDCS, whereas the stimulating area served as the moderator on the effect of TMS. Moderation analysis was conducted with the “meta” package in R.

### Publication bias analysis

Publication bias was evaluated by funnel plots and Egger’s regression intercept test.

Standard errors were plotted against the effect sizes in the funnel plots, with asymmetricity indicating publication bias (Copas & Shi, 2000). Egger’s regression test was based on the asymmetry of the funnel plot, which examined the relationship between outcomes and standard errors (Sterne & Egger, 2005). A significant result suggests a publication bias.

## Results

The available data was inadequate to perform a meta-analysis on several effects. Among the animal studies, there was only one article reported latency to freezing concerning the effect of anodal tDCS on long-term retention of contextual fear and cued fear (Manteghi et al., 2017). Additionally, there was only one article investigating the effect of cathodal tDCS on long-term fear retention (Duran et al., 2022). Therefore, the only available pooled effect size concerning long-term fear retention was the effect of anodal tDCS on contextual fear indexed by the percentage of freezing time. Moreover, there was inadequate data to perform a meta-analysis on the effect of TMS on the retention of fear among animal subjects. Among the three animal studies on the effect of TMS, two articles employed the contextual fear conditioning paradigm (Isogawa et al., 2003; Legrand et al., 2019), while the other used the cued fear conditioning paradigm (Baek et al., 2012). As for human studies, there was inadequate data to perform a meta-analysis on the effect of cathodal tDCS, with only two articles on fear extinction (Asthana et al., 2013; Ganho-Ávila et al., 2019) and one article on the return of fear (Ganho-Ávila et al., 2019). Although the available articles on these effects were not sufficient for running a meta-analysis, the main findings of the relevant articles were listed in Tables 1 and 2.

### Effect of tDCS on retention of fear among animal subjects

#### Effect of tDCS on short-term retention of contextual fear

Seven articles employed the facilitating anodal tDCS protocol, comprising 100 subjects in the active stimulation group and 70 subjects in the control or sham group. The pooled effect size of anodal tDCS on reducing the percentage of freezing time was -0.79 (*p* = .016, 95% CI [-1.42, -0.14]), while that on reducing the latency to freezing was 0.83 (*p* < .001, 95% CI [0.35, 1.31]) (Figures 2a i and 2a iii). The results suggested lower levels of fear retention in the active group compared to the sham group. In other words, anodal tDCS has a significant inhibiting effect on the short-term retention of contextual fear.

**Figure 2a.**
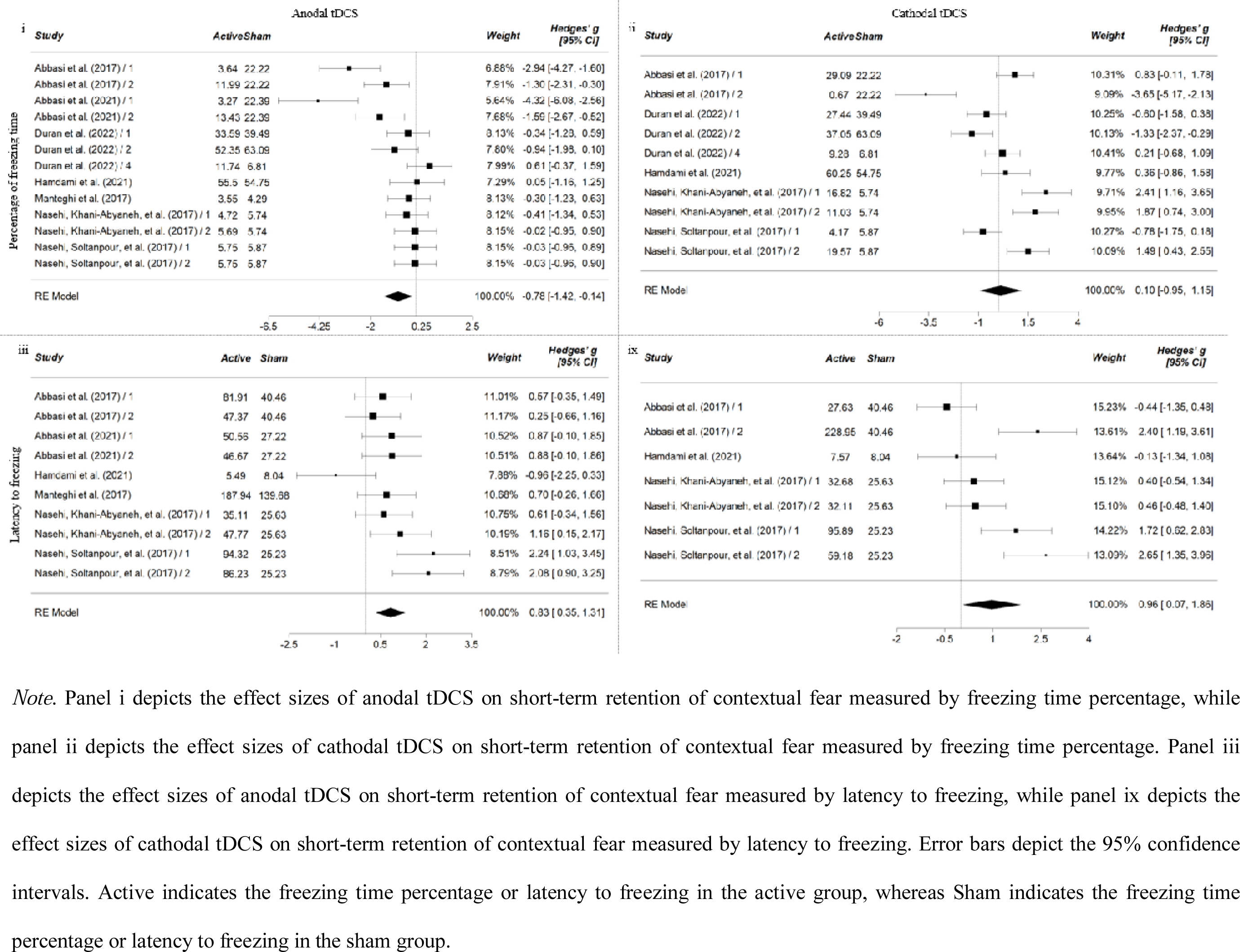
Forest Plot of Effect Sizes (Hedges’ g) of tDCS on Short-term retention of Contextual Fear among Animal Subjects.

Five articles employed the inhibiting cathodal tDCS protocol, comprising 74 subjects in the active stimulation group and 54 subjects in the control or sham group. The pooled effect size of cathodal tDCS on reducing the percentage of freezing time was 0.10 (*p* = .847, 95% CI [-0.95, 1.15]), while that on reducing the latency to freezing was 0.97 (*p* = .035, 95% CI [0.07, 1.86]) (Figures 2a ii and 2a ix). The results suggested the active group had lower levels of fear retention indexed by latency to freezing, compared to the sham group. In other words, cathodal tDCS has a significant inhibiting effect on the short-term retention of contextual fear measured by latency to freezing.

The results of the moderation analysis are illustrated in Figure S3. There was a significant moderating effect of stimulating duration on the effect of cathodal tDCS on short-term retention of contextual fear indexed by latency to freezing. Cathodal tDCS with a duration of 30 minutes has a stronger inhibiting effect on short-term retention of contextual fear, compared to 20-minute cathodal tDCS.

There was moderate to high heterogeneity across the anodal studies (*Q* = 46.56, *df* = 13, *p* < .001; *I^2^* = 79.7% for reducing freezing time percentage and *Q* = 19.49, *df* = 10, *p* = .021; *I^2^*= 53.91% for reducing latency to freezing, respectively) and cathodal studies (*Q* = 67.21, *df* = 10, *p* < .001; *I^2^*= 89.57% for reducing freezing time percentage and *Q* = 27.62, *df* = 7, *p* < .001; *I^2^* = 79.66% for reducing latency to freezing, respectively).

The funnel plot of the effect sizes of anodal tDCS on freezing time percentage showed asymmetric distribution (Figure S1a i). Moreover, Egger’s regression intercept test revealed a significant result (*z* = -4.996, *p* < .001), suggesting a publication bias. However, the publication bias on anodal studies reporting the latency to freezing was low, with symmetric distribution in the corresponding funnel plot (Figure S1a iii) and a non-significant result of Egger’s regression intercept test (*z* = 0.387, *p* = .699). As for the effect of cathodal tDCS, a publication bias among studies reporting latency to freezing was indicated by the asymmetric distribution in the funnel plot (Figure S1a ix) and a significant result (*z* = 2.315, *p* = .021) of Egger’s regression intercept test. However, the symmetric distribution in the funnel plot (Figure S1a ii) and a non-significant result (*z* = -0.986, *p* = .324) of Egger’s regression intercept test did not suggest a publication bias on cathodal studies reporting freezing time percentage.

#### Effect of tDCS on short-term retention of cued fear

Five articles employed the facilitating anodal tDCS protocol, comprising 61 subjects in the active stimulation group and 36 subjects in the control or sham group. The pooled effect size of anodal tDCS on reducing the percentage of freezing time was -0.33 (*p* = .116, 95% CI [-0.75, 0.08]), while that on reducing the latency to freezing was 0.45 (*p* = .017, 95% CI [0.08, 0.83]) (Figures 2b i and 2b iii). In other words, the active group reported lower levels of fear retention compared to the sham group. Anodal tDCS has an inhibiting effect on short-term retention of cued fear, which is significant when measured by the latency to freezing, but not significant when measured by the freezing time percentage.

**Figure 2b.**
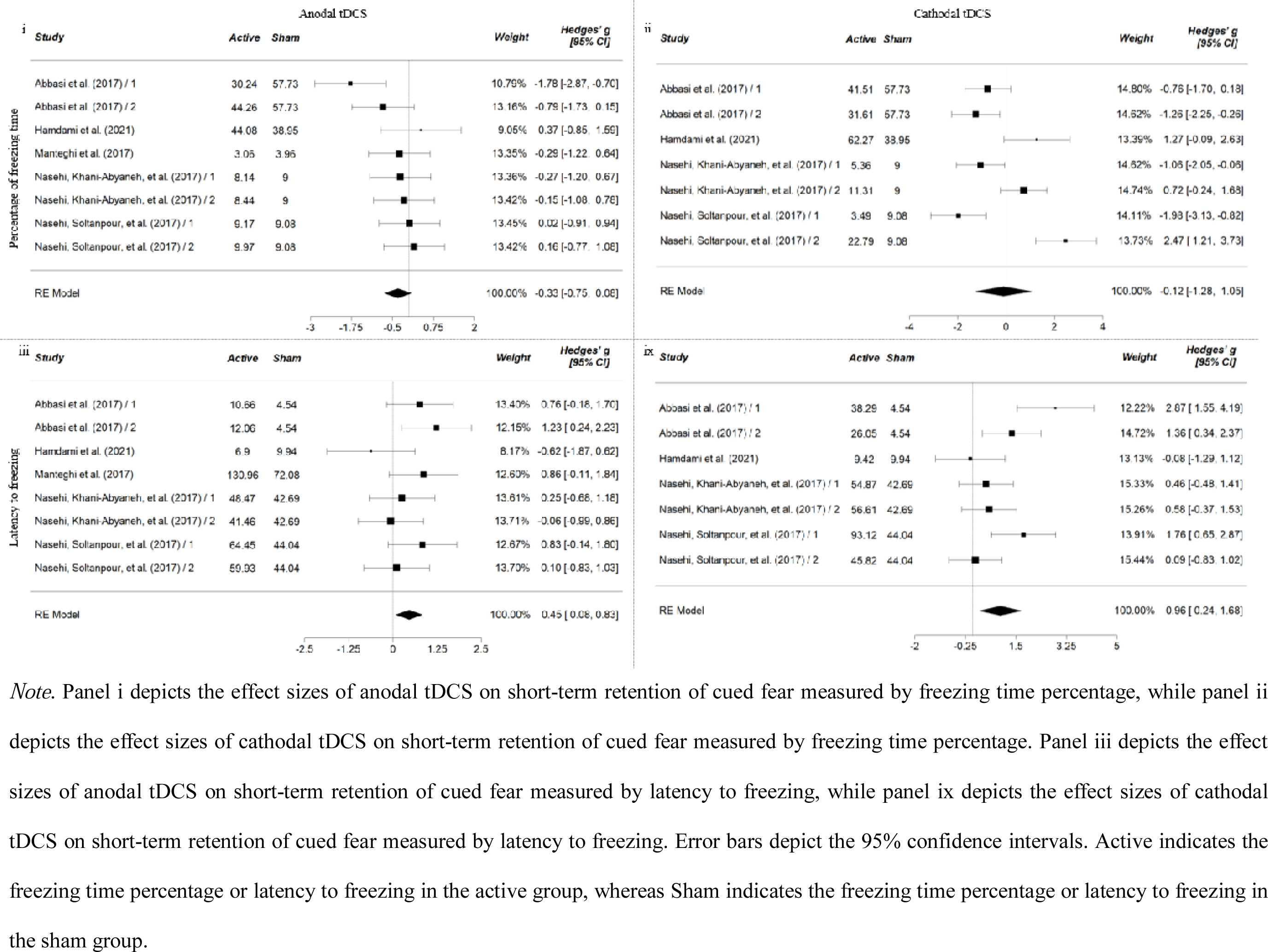
Forest Plot of Effect Sizes (Hedges’ g) of tDCS on Short-term retention of Cued Fear among Animal Subjects.

Four articles employed the inhibiting cathodal tDCS protocol, comprising 52 subjects in the active stimulation group and 28 subjects in the control or sham group. The pooled effect size of cathodal tDCS on reducing the percentage of freezing time was -0.12 (*p* = .846, 95% CI [-1.28, 1.05]), while that on reducing the latency to freezing was 0.96 (*p* = .009, 95% CI [0.24, 1.68]) (Figures 2b ii and 2b ix). In other words, the active group had lower levels of fear retention indexed by latency to freezing, but not freezing time percentage, compared to the sham group. Cathodal tDCS has a significant inhibiting effect on the short-term retention of contextual fear measured by latency to freezing.

The results of the moderation analysis are illustrated in Figure S3. There was a significant moderating effect of amplitude on the effect of anodal tDCS on short-term retention of cued fear indexed by latency to freezing. Anodal tDCS at 0.2 mA inhibits short-term retention of cued fear, whereas anodal tDCS at 2 mA enhance short-term retention of cued fear.

There was a moderate to high heterogeneity across both anodal studies (*Q* = 10.83, *df* = 8, *p* = .146; *I^2^* = 31.07% for reducing freezing time percentage and *Q* = 8.87, *df* = 8, *p* = .262; *I^2^*= 13.96% for reducing latency to freezing, respectively) and cathodal studies (*Q* = 42.81, *df* = 7, *p* < .001; *I^2^* = 87.74% for reducing freezing time percentage and *Q* = 18.34, *df* = 7, *p* = .005; *I^2^* = 69.68% for reducing latency to freezing, respectively).

Our findings indicated a significant publication bias in the studies of the anodal tDCS effect on short-term retention of cued fear. The funnel plot of the effect sizes of anodal tDCS showed symmetric distribution (Figure S1b i for freezing time percentage and Figure S1b iii for latency to freezing, respectively), and Egger’s regression intercept test revealed a non-significant result (*z* = -0.322, *p* = .748 for freezing time percentage and *z* = -1.028, *p* = .304 for latency to freezing, respectively). As for the effect of cathodal tDCS, the symmetric distribution in the funnel plot (Figure S1b ii) and a non-significant result (*z* = 1.8, *p* = .072) of Egger’s regression intercept test did not suggest a publication bias on cathodal studies reporting latency to freezing. While Egger’s regression intercept test revealed a non-significant result (*z* = 1.474, *p* = .141), the funnel plot showed asymmetric distribution (Figure S1b ix). Taken together, there was a potential publication bias on the effect of cathodal tDCS indexed by freezing time percentage.

#### Effect of tDCS on long-term retention of contextual fear

The pooled effect size of anodal tDCS on reducing the percentage of freezing time was 0.09 (*p* = .442, 95% CI [-0.14, 0.32]), indicating that anodal tDCS did not have a significant effect on long-term retention of contextual fear (Figure 2c i).

**Figure 2c.**
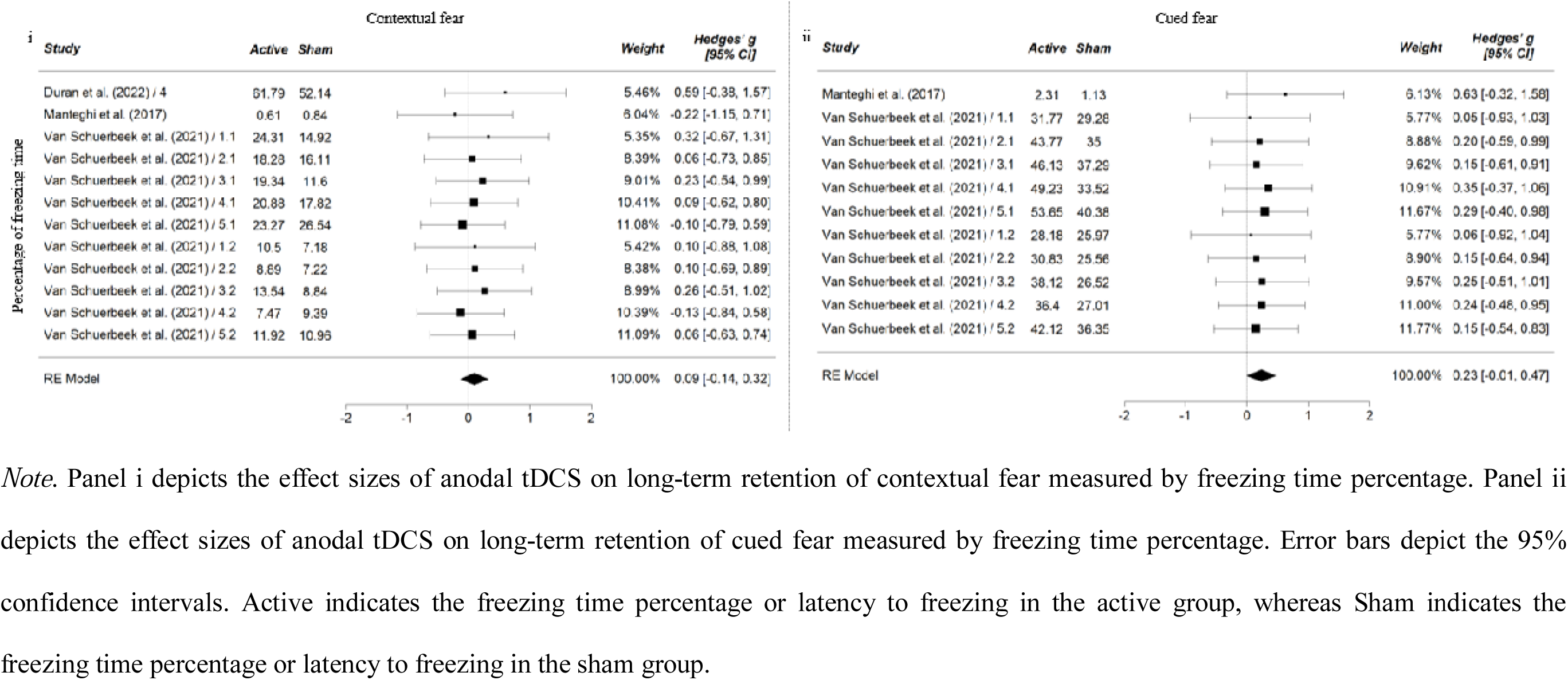
Forest Plot of Effect Sizes (Hedges’ g) of Anodal tDCS on Long-term retention of Fear among Animal Subjects.

The test of heterogeneity suggested homogeneity across anodal tDCS studies (*Q* = 2.65, *df* = 12, *p* = .995; *I^2^*= 0%). Additionally, tests did not reveal a significant publication bias. The funnel plot of the effect sizes of anodal tDCS showed symmetric distribution (Figure S1c i), and Egger’s regression intercept test revealed a non-significant result (*z* = 0.721, *p* = .471).

#### Effect of tDCS on long-term retention of cued fear

The pooled effect size of anodal tDCS on reducing the percentage of freezing time was 0.23 (*p* = .056, 95% CI [-0.01, 0.47]), indicating a marginally significant effect of anodal tDCS on long-term retention of cued fear (Figure 2c ii). The results suggested higher levels of fear retention in the active group than in the sham group.

The homogeneity across anodal tDCS studies (*Q* = 1.18, *df* = 11, *p* = 1; *I^2^* = 0%) was good, and there was no significant publication bias. The funnel plot of the effect sizes of anodal tDCS showed symmetric distribution (Figure S1c ii), and Egger’s regression intercept test revealed a non-significant result (*z* = -0.036, *p* = .971).

### Effect of tDCS on fear extinction among human subjects

Three articles employed the facilitating anodal tDCS protocol, comprising 113 participants in the active stimulation group and 68 participants in the control or sham group. The estimated pooled effect size of anodal tDCS on reducing the SCR difference during extinction was -0.55 (*p* = .050, 95% CI [-1.1, 0]), suggesting a significant facilitating effect of tDCS on fear extinction (Figure 3 i).

**Figure 3.**
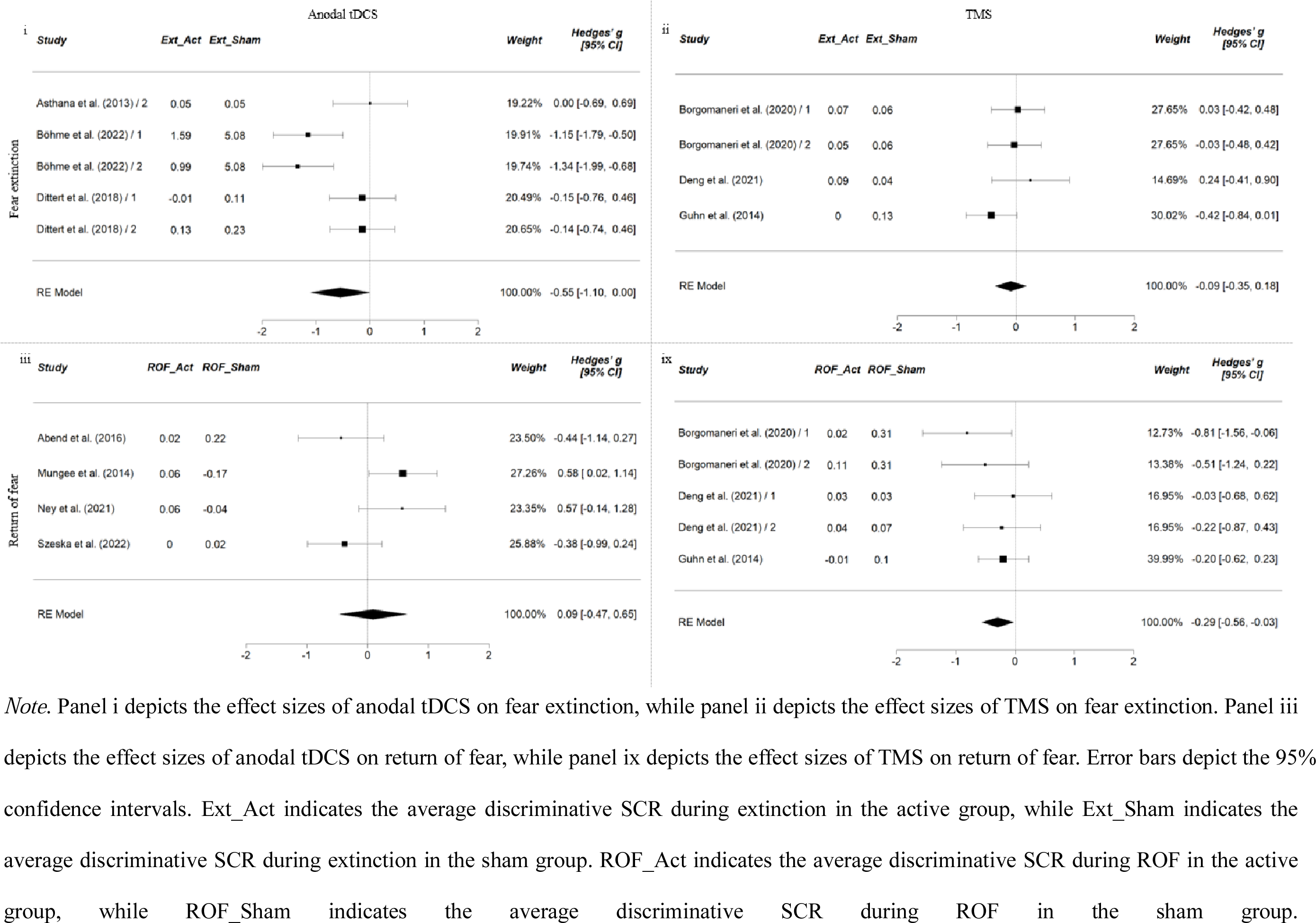
Forest Plot of Effect Sizes (Hedges’ g) of Human Studies

The results of the moderation analysis are illustrated in Figure S3. There was a significant moderating effect of amplitude on the effect of anodal tDCS on fear extinction. Anodal tDCS at 0.36 mA inhibits fear extinction, whereas such an inhibiting effect is weaker for anodal tDCS at 1.0 or 1.5 mA.

The test of heterogeneity suggested high heterogeneity across anodal tDCS studies (*Q* = 14.74, *df* = 5, *p* = .005; *I^2^* = 73.2%). While Egger’s regression intercept test revealed a non-significant result (*z* = -0.377, *p* = .706), the funnel plot showed asymmetric distribution (Figure S2 i). As such, there might be a potential publication bias on the effect of anodal tDCS on fear extinction.

### Effect of tDCS on return of fear among human subjects

Four articles employed the facilitating anodal tDCS protocol, comprising 78 participants in the active stimulation group and 72 participants in the control or sham group. The pooled effect size of anodal tDCS on reducing the discriminative SCR during ROF was 0.09 (*p* = .749, 95% CI [-0.47, 0.65]), indicating a non-significant facilitating effect of anodal tDCS on return of fear (Figure 3 iii).

There was significant heterogeneity across anodal studies (*Q* = 9.05, *df* = 4, *p* = .029; *I^2^* = 66.58%), but not a significant publication bias. The funnel plot of the effect sizes of anodal tDCS showed symmetric distribution (Figure S2 iii), and Egger’s regression intercept test revealed a non-significant result (*z* = -0.334, *p* = .738).

### Effect of TMS on fear extinction among human subjects

The pooled effect size of TMS on the reduction of the discriminative SCR was -0.09 (*p* = .511, 95% CI [-0.35, 0.18]), indicating a non-significant facilitating effect of TMS on fear extinction (Figure 3 ii). The test of heterogeneity suggests homogeneity across studies (*Q* = 3.59, *df* = 4, *p* = .309; *I^2^*= 18.15%). The funnel plot of the effect sizes of TMS showed symmetric distribution (Figure S2 ii), and Egger’s regression intercept test revealed a non-significant result (*z* = 1.3, *p* = .194). Taken together, these results did not suggest a significant publication bias.

### Effect of TMS on return of fear among human subjects

The pooled effect size of TMS on reducing the discriminative SCR was -0.29 (*p* = .032, 95% CI [-0.56, -0.03]), indicating a significant inhibiting effect of TMS on return of fear (Figure 3 ix). The moderating effect of the target area was not significant. There was neither a significant heterogeneity across studies (*Q* = 3.01, *df* = 5, *p* = .556; *I^2^* = 0%) nor a significant publication bias (Figure S2 ix; Egger’s regression intercept test: *z* = -0.954, *p* = .340).

## Discussion

The findings of the current meta-analytic study suggest that both anodal and cathodal tDCS of PFC inhibit short-term contextual and cued fear retrieval in animal studies (*Hedges’ g* = -0.79 for anodal effect on contextual fear indexed by freezing time percentage, *Hedges’ g* = 0.83 for anodal effect on contextual fear indexed by latency to freezing, *Hedges’ g* = 0.45 for anodal effect on cued fear indexed by latency to freezing, *Hedges’ g* = 0.97 for cathodal effect on contextual fear indexed by latency to freezing, *Hedges’ g* = 0.96 for cathodal effect on cued fear indexed by latency to freezing, respectively). For human studies, our findings suggested that anodal tDCS over the mPFC/vmPFC enhanced fear extinction (*Hedges’ g* = - 0.55), whereas TMS over the dlPFC and vmPFC reduced the return of fear (*Hedges’ g* = - 0.29).

### Effect of tDCS on fear retention in animal models

It was surprising in our review to find that anodal and cathodal tDCS stimulations exert similar inhibiting effects on short-term fear retention. Such a phenomenon was unexpected because stimulations with different polarities have hypothetically opposite effects on cortical excitability (Nitsche et al., 2008). A potential explanation is the divergent functions of subregions within the PFC. Specifically, the IL inhibits fear expression and prevents fear retention, whereas the prelimbic cortex promotes the return of fear (Milad & Quirk, 2002). Moreover, the anterior cingulate cortex (ACC) also plays a vital role in consolidating recent and remote fear memories (Einarsson & Nader, 2012; Frankland et al., 2004). Studies have revealed that the ACC enhances the recall and generalisation of fear (Einarsson et al., 2015; Sierra et al., 2017). The heterogeneous functions of subregions within the PFC pose difficulties for accurate neuromodulation by tDCS (Duran et al., 2022). Some studies have employed anodal tDCS to accelerate fear extinction, whereas others have employed cathodal tDCS to inhibit fear acquisition and consolidation.

Several aspects of tDCS may also account for the similar effects of anodal and cathodal stimulations on short-term fear retention. The relationship between stimulation and neural response is not only dependent on the stimulating polarity but also on the duration and intensity of tDCS, as well as the position of the reference electrode (Filmer et al., 2014). For example, Monte-Silva et al. (2013) reported that anodal tDCS could inhibit neural activation by increasing stimulation duration. Cathodal tDCS can also enhance cortical excitability with an increase in intensity (Batsikadze et al., 2013). Our moderation analysis results suggest that intensity exerts a significant moderating effect on the effect of anodal tDCS on the short-term retention of cued fear (indexed by latency to freezing, *z* = -3.076, *p* = .002), whereas the duration of stimulation moderated the effect of cathodal tDCS on the short-term retention of contextual fear (indexed by latency to freezing, *z* = 3.920, *p* < .001). Moreover, the tDCS effect may be associated with interference instead of the type of stimulation (Duran et al., 2022). For instance, distraction from the stimulation sensation was found to disrupt the consolidation of less persistent memories (Crestani et al., 2015).

Our findings reveal a moderate to large tDCS effect on the short-term retention of both contextual and cued fear. A common practice is to apply offline tDCS before threat acquisition. Cathodal tDCS with a duration of 30 minutes may be an effective protocol for inhibiting short-term fear retention, although there is less evidence to support the long-term effects of tDCS on the retention of fear.

### Effect of tDCS on threat extinction and return of fear in humans

Human studies have revealed anodal tDCS to exert a moderate and marginally significant effect on fear extinction. Although we did not detect any statistically significant moderators of tDCS in the human studies we reviewed, stimulation targeting the mPFC/vmPFC appears to enhance fear extinction (Böhme et al., 2022; Dittert et al., 2018; van ’t Wout et al., 2016; Vicario, Nitsche, et al., 2020), whereas stimulation over the dlPFC exerts no such effect (Asthana et al., 2013; Lipp et al., 2020). The vmPFC plays an essential role in the prefrontal-amygdala circuitry by integrating information from multiple inputs to exert top-down control on fear acquisition and extinction (Milad et al., 2007). Activation of the vmPFC may inhibit reactivity to aversive stimuli, thereby facilitating threat extinction.

Although there are inadequate data to perform a meta-analysis on the effect of cathodal tDCS, recent neuroimaging studies suggested that offline cathodal tDCS over the dlPFC prior to extinction can affect the neural responses to the CS during the extinction phase (Ganho-Ávila et al., 2022; Lee et al., 2023). Compared with the sham group in Ganho-Avila et al. (2022), the active group exhibited less activation in the prefrontal, postcentral, and paracentral regions when processing the CS+. The classification accuracy of the CS+ and CS-in the left anterior dorsal and ventral insulae and hippocampus was higher in the tDCS group than in the control group (Lee et al., 2023). Future studies might examine the effects of cathodal tDCS over the dlPFC to test the robustness of these findings.

In terms of the return of fear, our meta-analysis did not identify any significant effect of anodal tDCS on the return of fear in humans. The effect of cathodal tDCS on the return of fear is also unclear. Abend et al. (2016) found that stimulation over the right dlPFC before extinction enhanced the return of fear, whereas cathodal tDCS over the dlPFC inhibited the return of fear (Lipp et al., 2020; Mungee et al., 2016). The limited number of studies and inconsistent findings to date prohibit the identification of the effect of cathodal tDCS on the return of fear in human subjects. The null effect of tDCS on the return of fear is concerning from a clinical perspective because patients with anxiety disorders often suffer a relapse, and the return of fear is common in this patient population (Vervliet et al., 2013).

### Effect of TMS on threat extinction and return of fear in humans

Although our findings suggest that the effect of TMS is not apparent with respect to threat extinction, its inhibiting effect can be seen in studies measuring the return of fear. The pooled effect size (*Hedges’ g* = -0.29) was small but robust, as the effect was consistent across studies. This finding was also consistent with other TMS studies suggesting that TMS over the dlPFC and vmPFC reduces the return of fear (Raij et al., 2018; Oiala et al., 2022).

Owing to the small number of studies included in our review, we were unable to identify a specific TMS protocol that can be deemed more effective than others. However, there is some evidence suggesting that repetitive TMS over mPFC can facilitate fear extinction (Guhn et al., 2014). That continuous theta-burst TMS over the primary sensory cortex can lead to decreased fear retention (Ojala et al., 2022).

### Possible mechanisms of the effects of non-invasive brain stimulation

Our meta-analysis reveals that tDCS inhibits the short-term retention of both contextual and cued fear among animal subjects. A common practice is to apply offline tDCS over the frontal area right before threat acquisition. Such stimulation is likely to affect threat learning and/or consolidation, which is reflected in decreased retention of fear. Importantly, few animal studies focused on the effect of tDCS on fear extinction, which is a process of crucial clinal implications. Future studies might investigate whether and how anodal and cathodal tDCS affect fear extinction among animal subjects.

Human studies suggested an enhancing effect of anodal tDCS on fear extinction, as well as an inhibitory effect of TMS on return of fear. Timing of stimulation varied across studies. Brain stimulation immediately after threat acquisition is likely to inhibit the consolidation of fear memory (Asthana et al., 2013; Böhme et al., 2022; Borgomaneri et al., 2020). Furthermore, tDCS/TMS right before or during extinction training might contribute to the acquisition of the CS-no US association (Böhme et al., 2022; Dittert et al., 2018; Guhn et al., 2014), whereas stimulation right after extinction learning might enhance the consolidation of the newly learned non-fearful memory (Deng et al., 2021). Future studies might further investigate the mechanisms of the effects of non-invasive brain stimulation on threat extinction and the return of fear, which contribute to identifying the optimal timing of stimulation.

Few animal studies explored the effect of NIBS on fear extinction, which prevented us from conducting a meta-analysis of the relevant effects and comparing them with those among human subjects. There was only one study examining the effect of tDCS on fear extinction among animal subjects (Study 4 of Duran et al., 2022). The results suggested that cathodal tDCS right after extinction learning facilitates the consolidation of extinction memory. Additionally, Baek et al. (2012) found that online TMS during extinction enhances threat extinction and inhibits return of fear. The mechanism is likely to be that TMS contributes to the learning of the CS-no US association, which is consistent with human studies (Guhn et al., 2014; Raij et al., 2018). Future studies might further explore the effects of NIBS on fear extinction and return of fear among animal subjects. These results will be valuable for comparing the effects of NIBS and the relevant mechanisms across animal and human subjects, thus providing more reliable clinical implications.

### Implication for using non-invasive brain stimulation for anxiety disorders in humans

The treatment effect of tDCS on anxiety disorders in a previous review displays considerable heterogeneity in samples, anxiety measures, stimulation patterns, and outcomes, all of which prevented us from identifying the putative effect of tDCS (Stein et al., 2020). Fear conditioning model is a laboratory model of clinical anxiety that contributes to understanding the multi-faceted psychological and neurobiological mechanisms maladaptive of fear (Beckers et al., 2023). Our findings shed light on using NIBS for anxiety disorders in humans by reviewing the combination treatment effect of tDCS and threat extinction in the fear conditioning model. Overall, anodal tDCS over the mPFC/vmPFC enhanced fear extinction in this meta-analysis, which suggests that anodal tDCS is a promising tool to enhance the efficacy of exposure therapy. In the studies we reviewed, applying anodal tDCS over the mPFC/vmPFC with moderate intensity (e.g., 0.36 mA) appears to be an effective protocol.

In addition, our findings suggest that a combination of TMS and extinction learning has an inhibiting effect on the return of fear in humans. Specifically, repetitive TMS over dlPFC or mPFC before or after extinction learning may facilitate the new CS-no US associative learning, thereby reducing the return of fear. This finding is largely consistent with the existing literature that TMS has a robust treatment effect for treating PTSD and generalized anxiety disorder (Cirillo et al., 2019). While TMS may enhance the efficacy of exposure therapy, more research is needed to understand the neural pathways it targets and how it affects the existing threat association and/or the formation of the new non-threatening association. Neuroimaging methods might be adopted to investigate the underlying neural mechanisms.

### Limitations and future research directions

The limited number of studies in several areas prevented us from conducting a meta-analysis and identifying the relevant effects. Among animal subjects, for example, no meta-analysis could be performed to investigate the effect of cathodal tDCS or TMS on long-term fear retention. As for human studies, there were inadequate data to conduct a meta-analysis on the effect of cathodal tDCS on either fear extinction or the return of fear. More studies are required to further explore these effects.

Moreover, the heterogeneous designs of the selected studies made comparison difficult. For example, the studies we reviewed on the effect of tDCS on short-term fear retention among animal subjects employed divergent stimulation intensity, duration, and timing. As for the long-term retention of fear, the intervals between fear acquisition and long-term retention test varied from 15 days to 1 month. Among the studies involving human subjects, those investigating the effect of tDCS on fear extinction examined the stimulation of different brain regions. Future studies should employ more comparable stimulation protocols and fear conditioning paradigms to identify the optimal treatment.

A third limitation of our meta-analysis was the variance in effect sizes across studies and publication bias in several areas, including the effects of anodal and cathodal tDCS on short-term fear retention among animal subjects and the effect of anodal tDCS on fear extinction among human subjects. Future studies are advised to employ more comparable paradigms and practise pre-registration and open science to tackle the variance in effect sizes and publication bias.

## Conclusion

The findings of this meta-analytic study suggest that both anodal and cathodal tDCS of the PFC inhibit short-term fear retrieval in animal models. In human studies, anodal tDCS over the mPFC/vmPFC enhanced fear extinction, whereas TMS over the dlPFC and vmPFC has an inhibiting effect on the return of fear. The findings thus shed light on the optimal NIBS protocols for targeting the neural circuitry of threat extinction in humans.

## Author Contributions

Grace Lei and Charlene Lam conceived the study and developed the method. Grace Lei performed literature selection and data extraction. Grace Lei analyzed the data and wrote the manuscript. Cora Lai, Tatia Lee and Charlene Lam provided critical revisions. All authors approved the final version of the manuscript for submission.

