## Supplemental Material for "The Effect of Transcranial Direct Current and Magnetic Stimulation on Fear Extinction and Return of Fear: A meta-analysis and Systematic Review"

**Grace L.T. Lei^1,4^, Cora S.W. Lai^1,2^, Tatia M.C. Lee^1,3,5^, Charlene L.M. Lam*^1,4^**

^1^ The State Key Laboratory of Brain and Cognitive Sciences, The University of Hong Kong, Hong Kong, China.

^2^ School of Biomedical Sciences, LKS Faculty of Medicine, The University of Hong Kong, Pokfulam, Hong Kong

^3^ Laboratory of Neuropsychology and Human Neuroscience, The University of Hong Kong, Hong Kong, China.

^4^ Laboratory of Clinical Psychology and Affective Neuroscience, The University of Hong Kong, Hong Kong, China.

^5^ Guangdong-Hong Kong-Macao Greater Bay Area Center for Brain Science and Brain-Inspired Intelligence, Guangzhou, China

***Correspondence to:**

Charlene L.M. Lam, Ph.D.

Department of Psychology

Rm 6.59, 6/F, The Jockey Club Tower,

Centennial Campus, The University of Hong Kong

**Table S1a**

***Summary of the Human Studies on the effect of tDCS on Fear Conditioning Not Included in the Meta-analysis***

| Study | N (Groups) | N (total) | Gender M/F  (Mean Age ± SD) | Sample characteris-tics | Fear conditioning paradigm | CS / US | Number of Trials | tDCS current | tDCS duration | tDCS Montage | Number of tDCS sessions | Electrode position - cathode | Electrode position - anode | Timing of tDCS | Outcome | Other measure-  ments | Main Findings |
| --- | --- | --- | --- | --- | --- | --- | --- | --- | --- | --- | --- | --- | --- | --- | --- | --- | --- |
| Ganho-Ávila et al. (2022) | 16/18  (c-tDCS/  Sham) | 34 | 0/34  (23.32 ± 5.67) | Healthy female participants | Day 1 - acquisition  Day 2 - extinction | Colored squares / Scream | Acquisition: 12 CS+ (12 with US) / 16 CS-  Extinction: 10 CS+ / 10 CS- | 1 mA | 20 min | F4 | 1 | Right dlPFC (F4) | Contralateral deltoid | Offline (before extinction on Day 2) | fMRI, SCR, subjective ratings | ASI, STAI | C-tDCS decreased neural activity when processing CS + in prefrontal, postcentral and paracentral regions, during late extinction. |
| Lee et al. (2023) | 16/18  (c-tDCS/  Sham) | 34 | 0/34  (23.32 ± 5.67) | Healthy female participants | Day 1 - acquisition  Day 2 - extinction | Colored squares / Scream | Acquisition: 12 CS+ (12 with US) / 16 CS-  Extinction: 10 CS+ / 10 CS- | 1 mA | 20 min | F4 | 1 | Right dlPFC (F4) | Contralateral deltoid | Offline (before extinction on Day 2) | fMRI, SCR, subjective ratings | ASI, STAI | The classification accuracy of CS+ and CS- in the left anterior dorsal and ventral insulae and hippocampus was higher in the tDCS group than in the control group. |
| Lipp et al. (2020) | 20/20/20/20/20  (a-tDCS of cerebellum/  c-tDCS of cerebellum/  a-tDCS of dlPFC/  c-tDCS of dlPFC/  Sham) | 100 | 50/50  (24.51 ± 3.32) | Healthy participants | Day 1 - acquisition, extinction  Day 2 - extinction recall | Tone / Air puff | Acquisition: 96 paired CS-US  Extinction: 48 CS  Recall: 48 CS | Cere-  bellum: 2 mA  dlPFC:  1 mA | 20 min | Cere-  bellum: centered caudal  dlPFC: F3 | 1 | A-tDCS of cerebellum: right buccinator muscle  C-tDCS of cerebellum: centered caudal  A-tDCS of dlPFC: right supraorbital area  C-tDCS of dlPFC: F3 | A-tDCS of cerebellum: centered caudal  C-tDCS of cerebellum: right buccinator muscle  A-tDCS of dlPFC: F3  C-tDCS of dlPFC: right supraorbital area | Online (during extinction) | EMG | N.R. | A-tDCS of dlPFC inhibited fear consolidation. |
| Mungee et al. (2016) | 7/10  (c-tDCS/  Sham) | 17 | 5/12  (N.R.) | Healthy participants | Day 1 - acquisition  Day 2 - remind of CS+  Day 3 - fear assessment | Colored squares / Electric shock | Acquisition: 16 CS+ (6 with US) / 10 CS-  Remind of CS+: 1 CS+  Fear response assessment: 10 CS+ / 11 CS- | 1 mA | 20 min | F4 | 1 | right DLPFC (F4) | contralateral supraorbital | Offline (Day 2) | SCR | N.R. | No effect of c-tDCS was found. |
| Roesmann et al. (2022) | 27/26/26  (a-tDCS/  c-tDCS/  Sham) | 79 | A-tDCS: 12/15  (24.08 ± 3.14)  C-tDCS: 11/15  (24.89 ± 4.38)  Sham: 11/15  (24.32 ± 3.15) | Healthy participants | Acquisition, pre-stimulation test, post-stimulation test | Gabor patches / Fearful female face with scream | Acquisition: 60 CS+ (20 with US) / 60 CS-  Pre-stimulation test: 21 CS+ / 21 CS- / 21 each GS  Post-stimulation test: 21 CS+ / 21 CS- / 21 each GS | 1.5 mA | 10 min | anterior vmPFC | 1 | A-tDCS: under the chin  C-tDCS: on the forehead (anterior vmPFC) | A-tDCS: on the forehead (anterior vmPFC)  C-tDCS: under the chin | Offline (between pre-stimulation test and post-stimulation test) | MEG, pupil dilation, subjective ratings | CES-D,  LSAS, PANAS, SDS, SPAI, STAI, UI | C-tDCS inhibited fear generalization. |
| van ’t Wout et al. (2016) | 26/18  (Group 1/  Group 2) | 44 | Group 1: 16/10  (27.7 ± 8.45)  Group 2: 7/11  (26.72 ± 7.97) | Healthy participants | Day 1 - acquisition, extinction  Day 2 - extinction recall | Colored lights / Electric shock | Acquisition: 8 (5 with US) / 8 CS-  Extinction: 12 CS+ / 6 CS-  Extinction recall: 8 CS+ / 8 CS- | 2 mA | 10 min | AF3 | 1 | contralateral mastoid | vmPFC (AF3) | Group 1: Online (during 1^st^ extinction block)  Group 2: Online (during 2^nd^ extinction block) | SCR | N.R. | A-tDCS during 2^nd^ extinction block promoted late extinction learning compared with a-tDCS during 1^st^ extinction block. |
| van ’t Wout et al. (2017) | 14/14  (Group 1/  Group 2) | 28 | Group 1: 14/0  (53.36 ± 12.8)  Group 2: 14/0  (59.14 ± 11.1) | Males diagnosed with PTSD | Day 1 - acquisition, extinction  Day 2 - extinction recall | Colored lights / Electric shock | Acquisition: 8 CS+ (5 with US) / 8 CS-  Extinction: 6 CS+ / 6 CS-  Extinction recall: 8 CS+ / 8 CS- | 2 mA | 10 min | AF3 | 1 | contralateral mastoid | vmPFC (AF3) | Group 1: Online (during extinction)  Group 2: Offline (after extinction on Day 1) | SCR | BAI, BDI-II, Hoge Combat Scale | A-tDCS during extinction promoted early extinction recall compared with a-tDCS after extinction. |
| Vicario, Nitsche, et al. (2020) | 16/16  (a-tDCS/  Sham) | 32 | tDCS: 5/11  (23.93 ± 5.26)  Sham: 5/11  (24.37 ± 4.58) | Healthy participants | Day 1 - acquisition, extinction  Day 2 - extinction recall | Colored circles / Electric shock | Acquisition: 7 CS+ (5 with US) / 7 CS-  Extinction: 10 CS+ / 10 CS-  Extinction recall: 10 CS+ / 10 CS- | 2 mA | 10 min | AF3 | 1 | contralateral mastoid | vmPFC (AF3) | Online (during extinction) | SCR, subjective ratings | DASS, AUDIT | A-tDCS facilitated extinction learning and recall. |

*Note*. A-tDCS refers to anodal tDCS, while c-tDCS refers to cathodal tDCS. ASI indicates Anxiety Sensitivity Index (Reiss et al., 1986). AUDIT indicates Alcohol Use Disorders Identification Test (Saunders et al., 1993). BAI indicates Beck Anxiety Inventory (Beck et al., 1993). BDI indicates Beck Depression Inventory (Beck et al., 1996). CES-D indicates Center for Epidemiologic Studies Depression Scale (Sheehan et al., 1995). DASS indicates Depression Anxiety Stress Scales (Lovibond & Lovibond, 1995). LSAS indicates Liebowitz Social Anxiety Scale (Liebowitz, 1987). PANAS indicates Positive and Negative Affect Schedule (Watson et al., 1988). SDS indicates Social Desirability Scale (Crowne & Marlowe, 1960). SPAI indicates Social Phobia and Anxiety Inventory (Turner et al., 1989). STAI indicates State-Trait Anxiety Inventory (Spielberger, 1983). UI indicates Intolerance of Uncertainty scale (Buhr & Dugas, 2002). EMG refers to electromyography. fMRI refers to functional magnetic resonance imaging. MEG refers to magnetoencephalography. N.R. refers to not reported in the article.

**Table S1b**

***Summary of the Human Studies on TMS and Fear Conditioning Not Included in the Meta-analysis***

| Study | N (Groups) | N (total) | Gender M/F  (Mean Age ± SD) | Sample characteristics | Fear conditioning Paradigm | CS / US | Number of Trials | TMS target | Timing of TMS | Outcome | Other measurements | Main Findings |
| --- | --- | --- | --- | --- | --- | --- | --- | --- | --- | --- | --- | --- |
| Ojala et al. (2022) | 28/34  (left TMS/  right Sham) | 68 | 34/34  (23.7 ± 4.2) | Healthy participants | Day 1 - acquisition  Day 2 - recall test, relearning | 4 s electric pulses to the left hand / 500 ms electric shock to the foot | Acquisition: 48 CS+ (24 with US) / 48 CS-  Recall test: 12 CS+ / 12 CS-  Relearning: 48 CS+ (24 with US) / 48 CS- | S1 | Offline (before acquisition on Day 1) | Fear-potentiated startle, pupil dilation, SCR, subjective ratings | N.R. | cTBS inhibited return of fear. |
| Raij et al. (2018) | 28 | 28 | 23/5 (28 ± N.R.) | Healthy participants | Day 1 - acquisition  Day 2 - extinction learning  Day 3 - extinction recall | Colored lights / Electric shock | Acquisition: 8 CS+ (5 with US) / 4 CS-  Extinction learning: 8 CS+ / 4 CS-  Extinction recall: 8 CS+ / 4 CS- | vmPFC | Online (during extinction learning on Day 2) | SCR | N.R. | Online TMS on vmPFC inhibited return of fear. |

*Note*. cTBS refers to continuous theta-burst transcranial magnetic stimulation. S1 refers to the primary sensory cortex. N.R. refers to not reported in the article.

**Figure S1a**

***Funnel Plot of Effect Sizes of tDCS on Short-term retention of Contextual Fear among Animal Subjects***

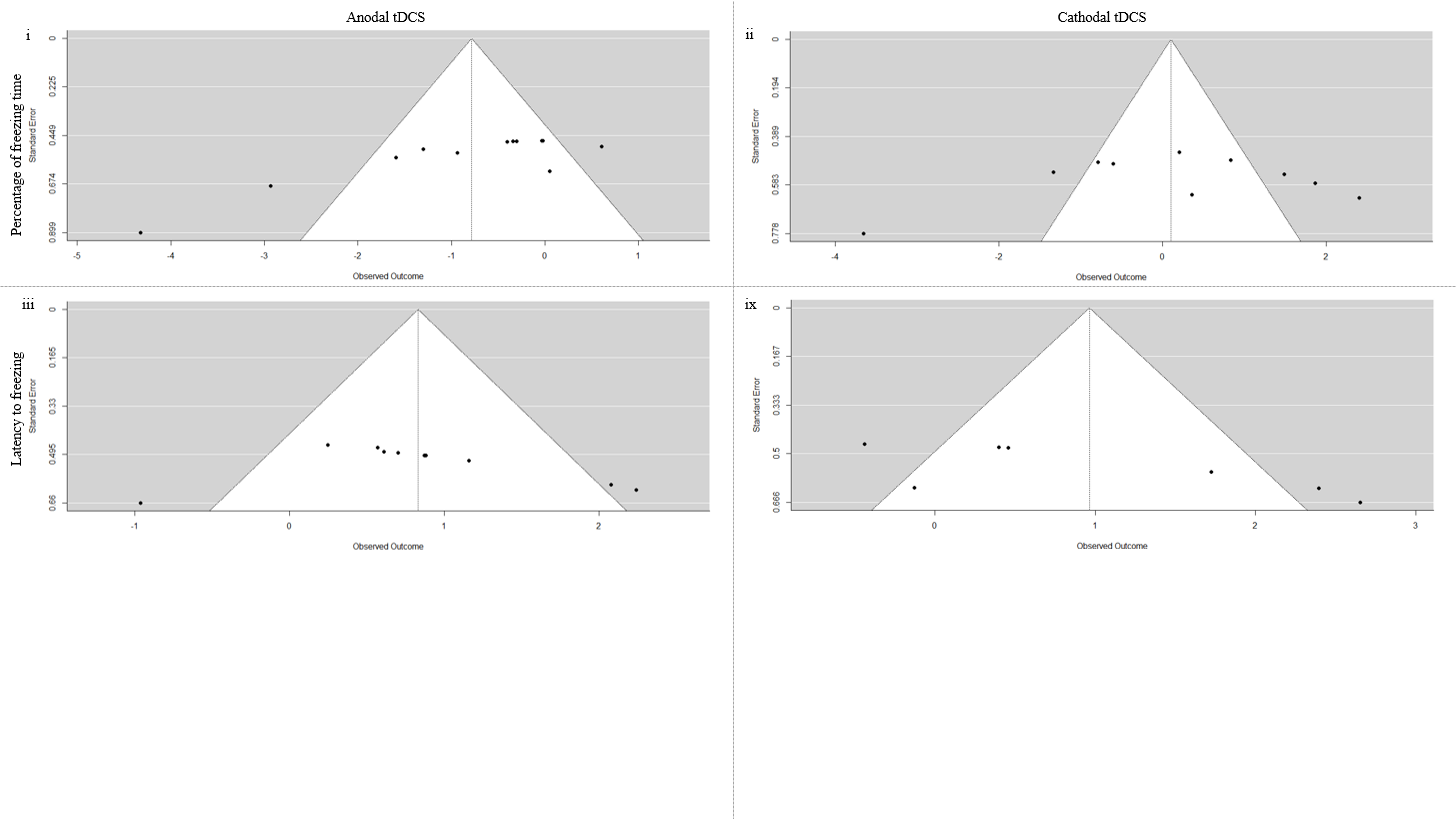

*Note*. Panel i depicts the effect sizes of anodal tDCS on short-term retention of contextual fear measured by freezing time percentage, while panel ii depicts the effect sizes of cathodal tDCS on short-term retention of contextual fear measured by freezing time percentage. Panel iii depicts the effect sizes of anodal tDCS on short-term retention of contextual fear measured by latency to freezing, while panel ix depicts the effect sizes of cathodal tDCS on short-term retention of contextual fear measured by latency to freezing.

**Figure S1b**

***Funnel Plot of Effect Sizes of tDCS on Short-term retention of Cued Fear among Animal Subjects***

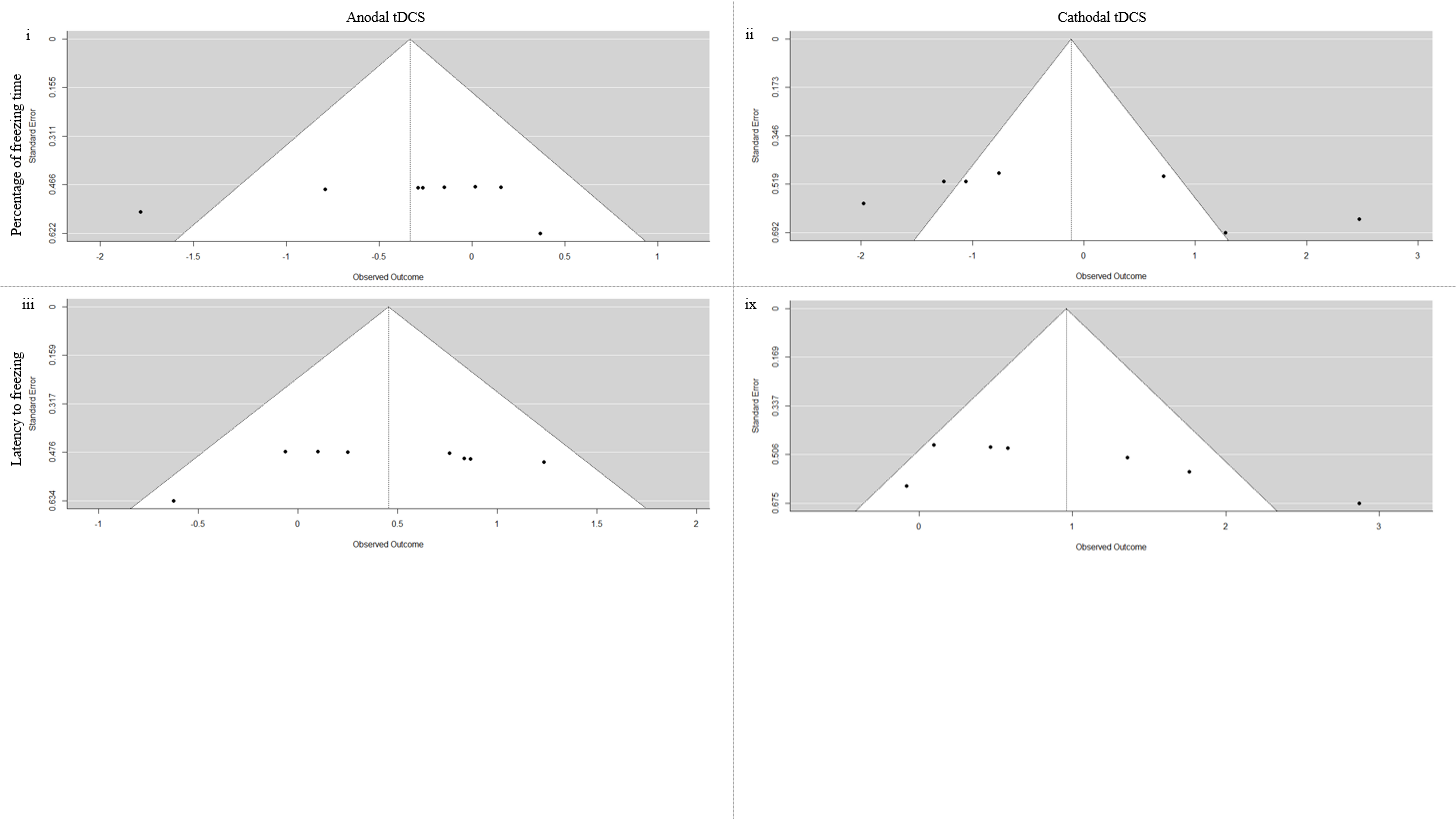

*Note*. Panel i depicts the effect sizes of anodal tDCS on short-term retention of cued fear measured by freezing time percentage, while panel ii depicts the effect sizes of cathodal tDCS on short-term retention of cued fear measured by freezing time percentage. Panel iii depicts the effect sizes of anodal tDCS on short-term retention of cued fear measured by latency to freezing, while panel ix depicts the effect sizes of cathodal tDCS on short-term retention of cued fear measured by latency to freezing.

**Figure S1c**

***Funnel Plot of Effect Sizes of Anodal tDCS on Long-term retention of Fear among Animal Subjects***

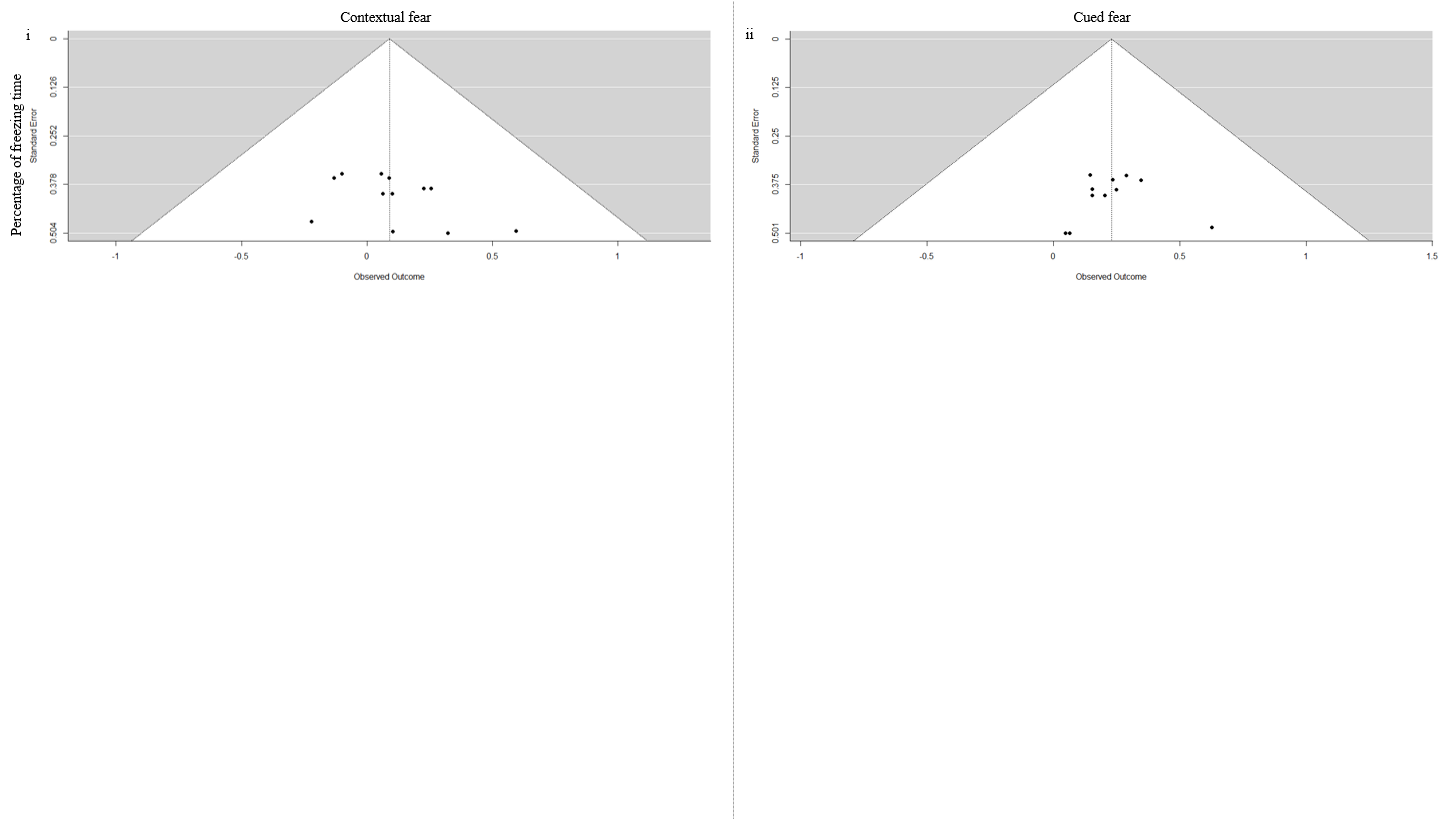

*Note*. Panel e depicts the effect sizes of anodal tDCS on long-term retention of contextual fear measured by freezing time percentage. Panel f depicts the effect sizes of anodal tDCS on long-term retention of cued fear measured by freezing time percentage.

**Figure S2**

***Funnel Plot of Effect Sizes of Human Studies***

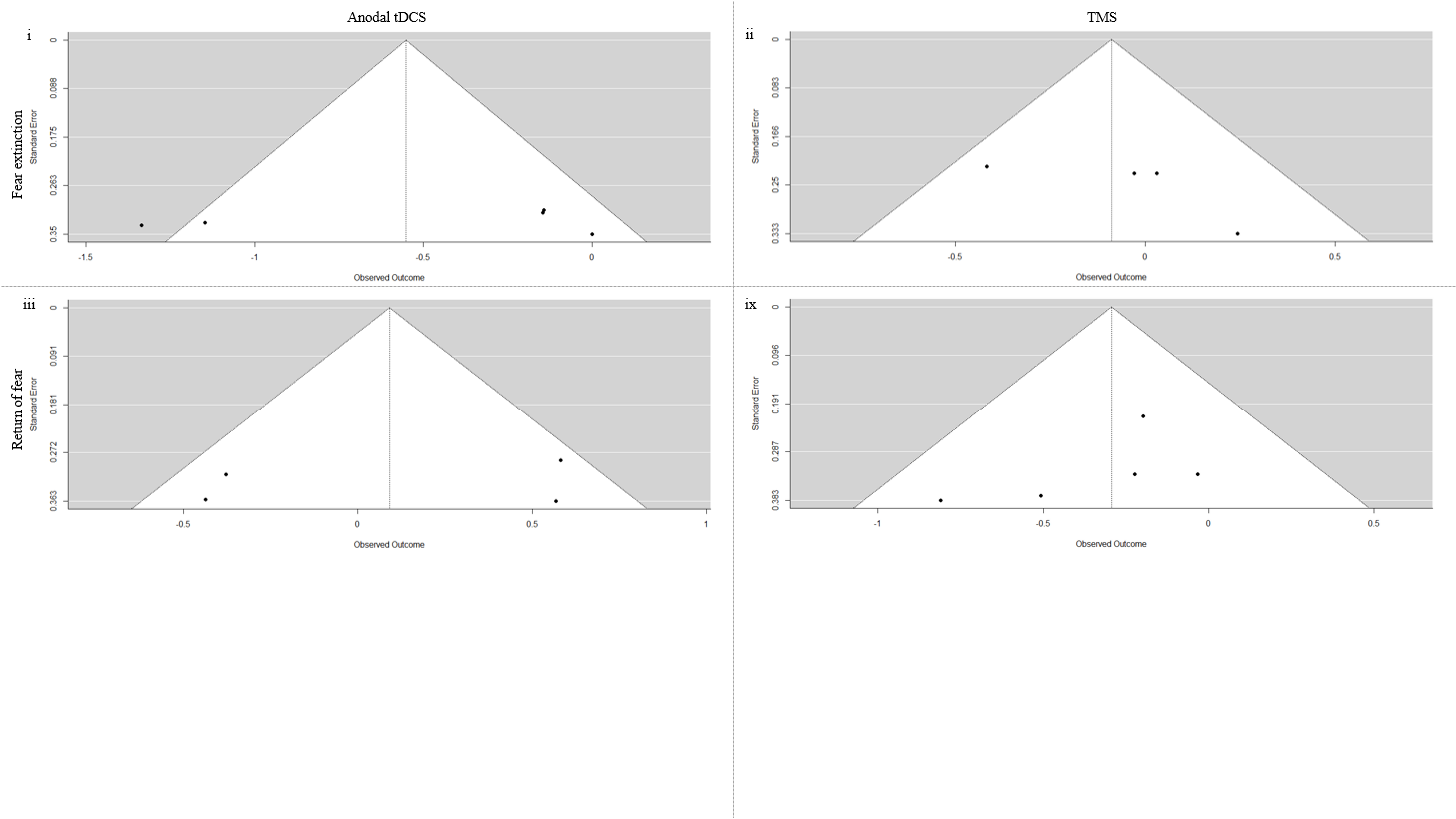

*Note*. Panel i depicts the effect sizes of anodal tDCS on fear extinction, while panel ii depicts the effect sizes of TMS on fear extinction. Panel iii depicts the effect sizes of anodal tDCS on return of fear, while panel ix depicts the effect sizes of TMS on return of fear.

**Figure S3**

***Results of Moderation Analysis***

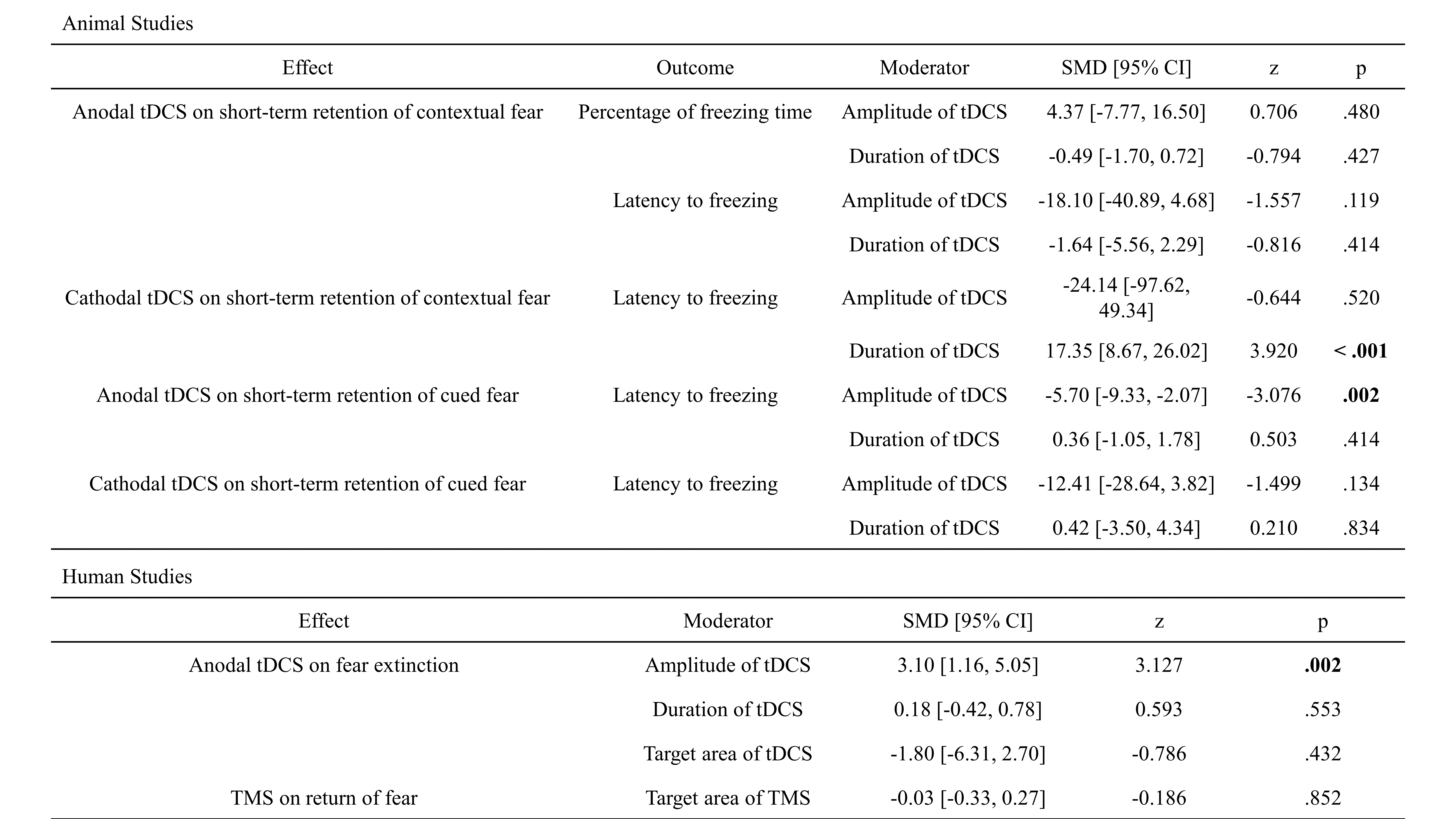
